# MprF-mediated immune evasion is necessary for *Lactiplantibacillus plantarum* resilience in *Drosophila* gut during inflammation

**DOI:** 10.1101/2024.01.09.574795

**Authors:** Aranzazu Arias-Rojas, Adini Arifah, Georgia Angelidou, Belal Alshaar, Ursula Schombel, Emma Forest, Dagmar Frahm, Volker Brinkmann, Nicole Paczia, Chase Beisel, Nicolas Gisch, Igor Iatsenko

## Abstract

**Background:** Multiple peptide resistance factor (MprF) confers resistance to cationic antimicrobial peptides (AMPs) in several pathogens, thereby enabling evasion of the host immune response. While MprF has been proven to be crucial for the virulence of various pathogens, its role in commensal gut bacteria remains uncharacterized. To close this knowledge gap, we used a common gut commensal of animals, *Lactiplantibacillus plantarum*, and its natural host, the fruit fly *Drosophila melanogaster*, as an experimental model to investigate the role of MprF in commensal-host interactions.

**Results:** The *L. plantarum* Δ*mprF* mutant that we generated exhibited deficiency in the synthesis of lysyl-phosphatidylglycerol (Lys-PG), resulting in increased negative cell surface charge and increased susceptibility to AMPs. Susceptibility to AMPs had no effect on Δ*mprF* mutant’s ability to colonize guts of uninfected flies. However, we observed significantly reduced abundance of the Δ*mprF* mutant after infection-induced inflammation in the guts of wild-type flies but not flies lacking AMPs. These results demonstrate that host AMPs reduce the abundance of the Δ*mprF* mutant during infection. We found in addition that the Δ*mprF* mutant compared to wild-type *L. plantarum* induces a stronger intestinal immune response in flies due to the increased release of immunostimulatory peptidoglycan fragments, indicating an important role of MprF in promoting host tolerance to commensals.

**Conclusion:** Overall, our results demonstrate that MprF, besides its well-characterized role in pathogen immune evasion and virulence, is also an important resilience factor in maintaining stable microbiota-host interactions during intestinal inflammation.

## Introduction

Gut microbial communities exist in an open ecosystem where they are subject to various perturbations, like exposure to toxins, dietary changes, and infections [1]. Inflammatory responses induced by infections are among the most frequent disruptions that gut-associated microbial communities experience over the lifespan of an individual [2]. During an intestinal inflammatory response, numerous antimicrobial effectors are produced to suppress and eliminate pathogens [3]. These immune effectors are often non-specific and target conserved molecular patterns present in both pathogenic and commensal bacteria, yet healthy gut microbiota can remain stable for decades in humans [4]. Hence, gut commensals exhibit resilience to gut intestinal immune responses [5, 6]. As one important example, human commensals from the phylum Bacteroidetes alter their lipopolysaccharide structure, which enhances resistance to antimicrobial peptides (AMPs) and facilitates commensal resilience during gut inflammation [7]. Iron limitation induced by infection is another host defence reaction that non-specifically targets gut commensals and restricts their access to this essential nutrient [8–11]. *Bacteroides thetaiotaomicron* was shown to acquires iron through siderophores produced by the other gut bacteria [12]. Such siderophore cross-feeding between different bacteria allows gut commensals to acquire iron in the inflamed gut and promotes gut microbiota resilience. However, with the exception of these few studies and despite the importance for host health, microbiota resilience mechanisms to inflammation remain little studied.

The fruit fly *Drosophila melanogaster* has been widely used as a genetically-tractable model to study host–microbe interactions, including microbiota resilience mechanisms [13–19]. Fruit flies rely on cellular and humoral arms of defence against invading pathogens [20–22]. Two major cell types of hemocytes function in cellular immune defence: plasmatocytes – involved in phagocytosis, and crystal cells which mediate the melanisation reaction [23, 24]. This reaction is particularly important against *S. aureus* infection [25].

Infection-induced synthesis and secretion of AMPs is the hallmark of *Drosophila* humoral immune response [26]. This response is regulated mainly by two conserved NF-κB pathways: Toll and Imd. The Toll pathway can be activated by bacterial proteases, fungal glycans or Lysine-type peptidoglycan (PGN) from Gram-positive bacteria [27]. Extracellular receptors PGRP-SA and GNBP1 form a complex that recognizes Lys-type PGN, ultimately activating the Toll signalling cascade and synthesis of antimicrobial effectors [28]. The Imd pathway is initiated by diaminopimelic (DAP)-type PGN sensed by transmembrane receptor PGRP-LC or by intracellular receptor PGRP-LE, resulting in nuclear translocation of NF-κB transcription factor Relish and expression of AMPs [29, 30]. While both Toll and Imd pathways regulate a systemic immune response, only Imd controls intestinal AMP expression [31].

Flies lacking major AMP classes were recently generated and proved to be instrumental in demonstrating an essential role of AMPs in vivo in the defence against Gram-negative pathogens and in the control of gut microbiota [32–34]. Given that the majority of the *Drosophila* microbiota members produce DAP-type PGN, they elicit Imd pathway activation in the gut [35–38]. However, in contrast to pathogens, commensals induce a mild AMP response due to the tolerance mechanisms deployed by the host [39]. One such tolerance mechanism is the potent induction of multiple negative regulators that fine-tune Imd pathway activation at different levels. These negative regulators are necessary to maintain host-microbiota homeostasis by preventing chronic deleterious Imd pathway activation and potent AMP response that would target gut commensals [40–43].

Recently, we demonstrated that, besides host tolerance to microbiota, commensal-encoded resilience mechanisms are essential to maintain stable microbiota-host associations, particularly during intestinal inflammation [14]. Specifically, we showed that *Drosophila* microbiota composition and abundance remain stable during infection. Using the prominent *Drosophila* commensal *Lactiplantibacillus plantarum* as a model, we demonstrated that resistance to AMPs is an essential commensal resilience mechanism during intestinal inflammation. We identified the *dlt* operon involved in the esterification of teichoic acids with D-alanine as one of the mediators of *Lp* resistance to *Drosophila* AMPs [14]. Considering that the *dlt* operon is also an important virulence factor of several pathogens that protect them from host AMPs [44, 45], our work illustrated that mechanisms typically associated with virulence can also be exploited by commensals to maintain association with the host.

Here, we explored the generality of this phenomenon and investigated the role of additional genes associated with pathogen AMP resistance in the commensal resilience during inflammation. We focused on the multiple peptide resistance factor (MprF) protein, which confers resistance to AMPs in several bacteria [46]. MprF is an integral membrane enzyme that catalyzes the alteration of the negatively-charged lipid phosphatidylglycerol (PG) with L-lysine, thereby neutralizing the membrane surface charge and providing resistance to AMPs [47, 48]. The resulting modified lipid, lysyl-phosphatidylglycerol (Lys-PG), is produced by MprF using phosphatidylglycerol and aminoacyl-tRNA as substrates [49–51]. Since Δ*mprF* mutants originally identified in *S. aureus* were susceptible to multiple AMPs due to the lack of Lys-PG, the gene was named *multiple peptide resistance factor* [52]. After that, deficiency in Lys-PG synthesis in Δ*mprF* mutants was linked to cationic AMP susceptibility in several other bacteria, including *Bacillus anthracis*, *B. subtilis*, *Enterococcus faecalis*, *Listeria monocytogenes*, *Mycobacterium tuberculosis*, and *Pseudomonas aeruginosa* [53–59]. MprF proteins proved to be crucial for the virulence of various pathogens, thereby demonstrating an essential role of MprF in bacterial immune evasion and making it an attractive target for the development of antivirulence strategies [60].

While the function of MprF in immune evasion and antibiotic resistance of pathogens has been extensively studied, the role of MprF in commensal bacteria remains uncharacterized.

Here, we used the *Drosophila* commensal *L. plantarum* as a model to investigate the involvement of MprF in commensal-host interactions. Using a newly generated *L. plantarum* Δ*mprF* mutant, we showed that it is impaired in the synthesis of Lys-PG, leading to increased negative cell surface charge and increased susceptibility to several AMPs. Consequently, the abundance of the Δ*mprF* mutant in the *Drosophila* gut was significantly reduced after infection- or genetically-induced inflammation. Hence, our results demonstrate an essential role of MprF-mediated AMP resistance in commensal resilience during inflammation.

### *mprF* is required for *S. aureus* virulence and resistance to *Drosophila* AMPs

Before attempting to generate *mprF*-deficient fruit fly commensals, we first wanted to prove with existing *mprF* mutants the relevance of *mprF* in *Drosophila* model. We decided to use the existing *S. aureus* Δ*mprF* mutant to test whether *mprF* is required for pathogen virulence in *Drosophila* model due to increased sensitivity to host AMPs, as was shown in other animal models [52]. For this purpose, we performed systemic infections of fruit flies by needle pricking with an *S. aureus* wild-type and an isogenic Δ*mprF* mutant. By monitoring the survival of infected flies, we found that the Δ*mprF* mutant is significantly attenuated compared to wild-type *S. aureus* (Fig. 1a and Fig. S1a) in flies of four different genetic backgrounds. Additionally, we estimated pathogen growth within the host by quantifying bacterial CFUs in fly homogenates. Consistent with survival, wild-type flies efficiently controlled Δ*mprF* mutant growth, as bacterial numbers didn’t increase significantly over the course of infection. In contrast, wild-type *S. aureus* reached significantly higher load compared to the Δ*mprF* mutant, especially at 21 hours post-infection. (Fig. 2b).

**Figure 1.**
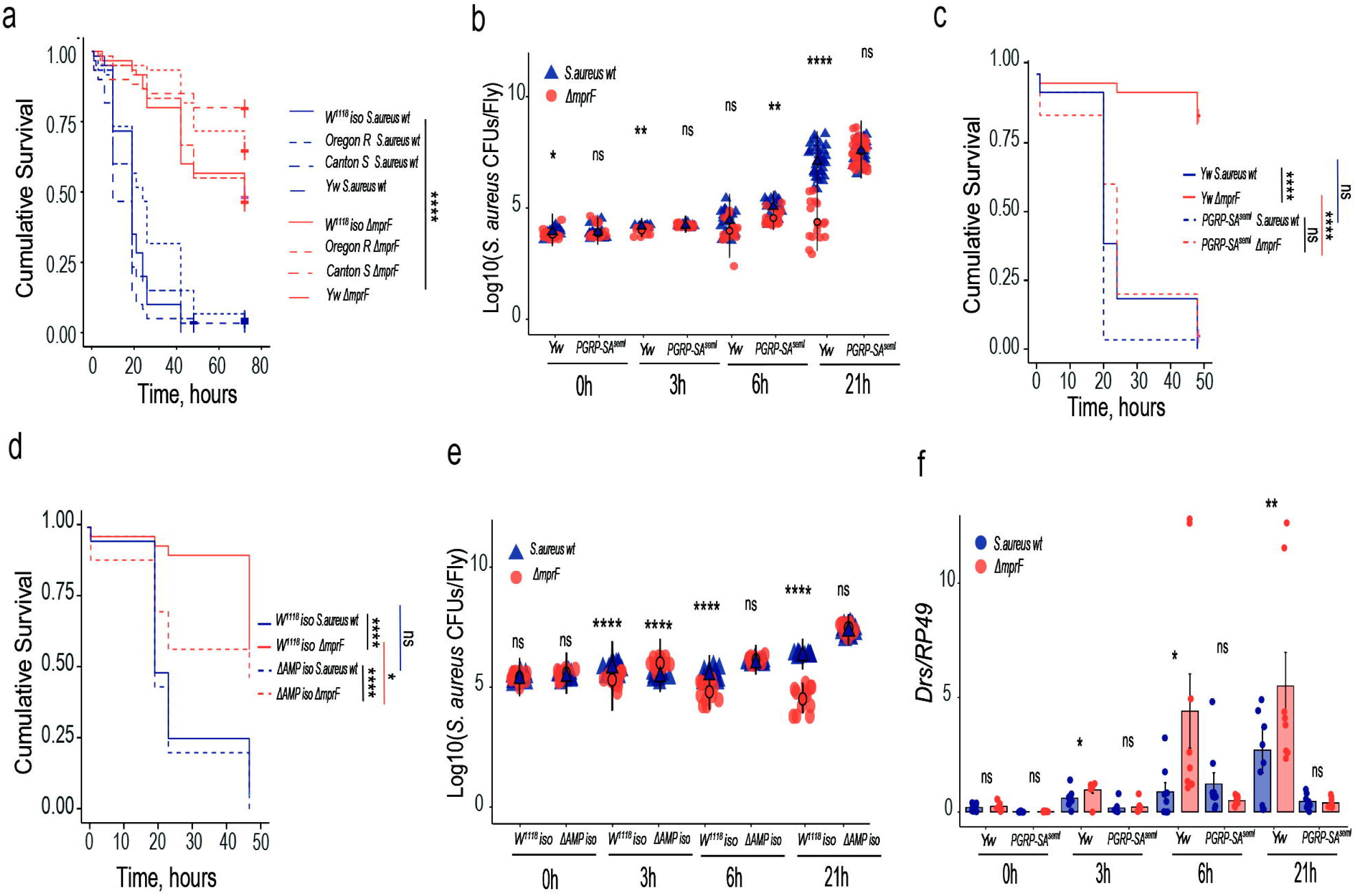
MprF is required for *S. aureus* virulence and resistance to *Drosophila* AMPs. (**a**) Survival rates of *Drosophila* wild-type strains infected with wild-type *S. aureus* or *S. aureus* Δ*mprF* mutant (n=3, independent experiments). (**b**) Measurement of *S. aureus* wild-type (n1) or *S. aureus* Δ*mprF* burden (n2) in wild-type (*yw*) and *PGRP-SA^seml^* flies. Number of samples (n) in *yw* at 0h (n1=17, n2=18), 3h (n1=12, n2=12), 6h (n1=18, n2=16) and 21h (n1=32, n2=17). *PGRP-SA^seml^* at 0h (n1=19, n2=20), 3h (n1=12, n2=12), 6h (n1=16, n2=20) and 21h (n1=27, n2=33). (**c**) Survival rates of *yw* and *PGRP-SA^seml^* flies infected with *S. aureus* wild-type or *S. aureus* Δ*mprF* mutant. (**d**) Survival rates of *w^1118^ iso* and Δ*AMP* flies infected with *S. aureus* wild-type or *S. aureus* Δ*mprF* mutant. (**e**) *S. aureus* wild-type (n1) and *S. aureus* Δ*mprF* mutant (n2) load in *w^1118^ iso* and Δ*AMPs* flies. Number of samples (n) in *w^1118^ iso* at 0h (n1=12, n2=12), 3h (n1=12, n2=12), 6h (n1=12, n2=12) and 21h (n1=12, n2=12). Δ*AMPs* at 0h (n1=12, n2=12), 3h (n1=12, n2=12), 6h (n1=12, n2=12) and 21h (n1=12, n2=12). (**f**) *Drosomycin (Drs)* gene expression in *yw* and *PGRP-SA^seml^* flies infected with *S. aureus* wild-type (n1) and *S. aureus* Δ*mprF* mutant (n2). Number of samples (n) in *yw* at 0h (n1=9, n2=9), 3h (n1=8, n2=8), 6h (n1=8, n2=9) and 21h (n1=9, n2=8). *PGRP-SA^seml^* at 0h (n1=8, n2=9), 3h (n1=8, n2=8), 6h (n1=9, n2=9) and 21h (n1=8, n2=8). Each sample contains 5 animals. Single dots in the bar plot show gene expression from pools of n=5 animals. Single dots are mean CFU values from pools of n=5 animals in the log10 scale. Black rounded dots show the median. Whiskers show either lower or upper quartiles. Each survival graph shows cumulative results of three independent experiments. In all figures, *p < 0.05, **p < 0.01, ***p < 0.001. Kruskal-Wallis and Bonferroni post hoc tests were used for the statistical analysis.

**Figure 2.**
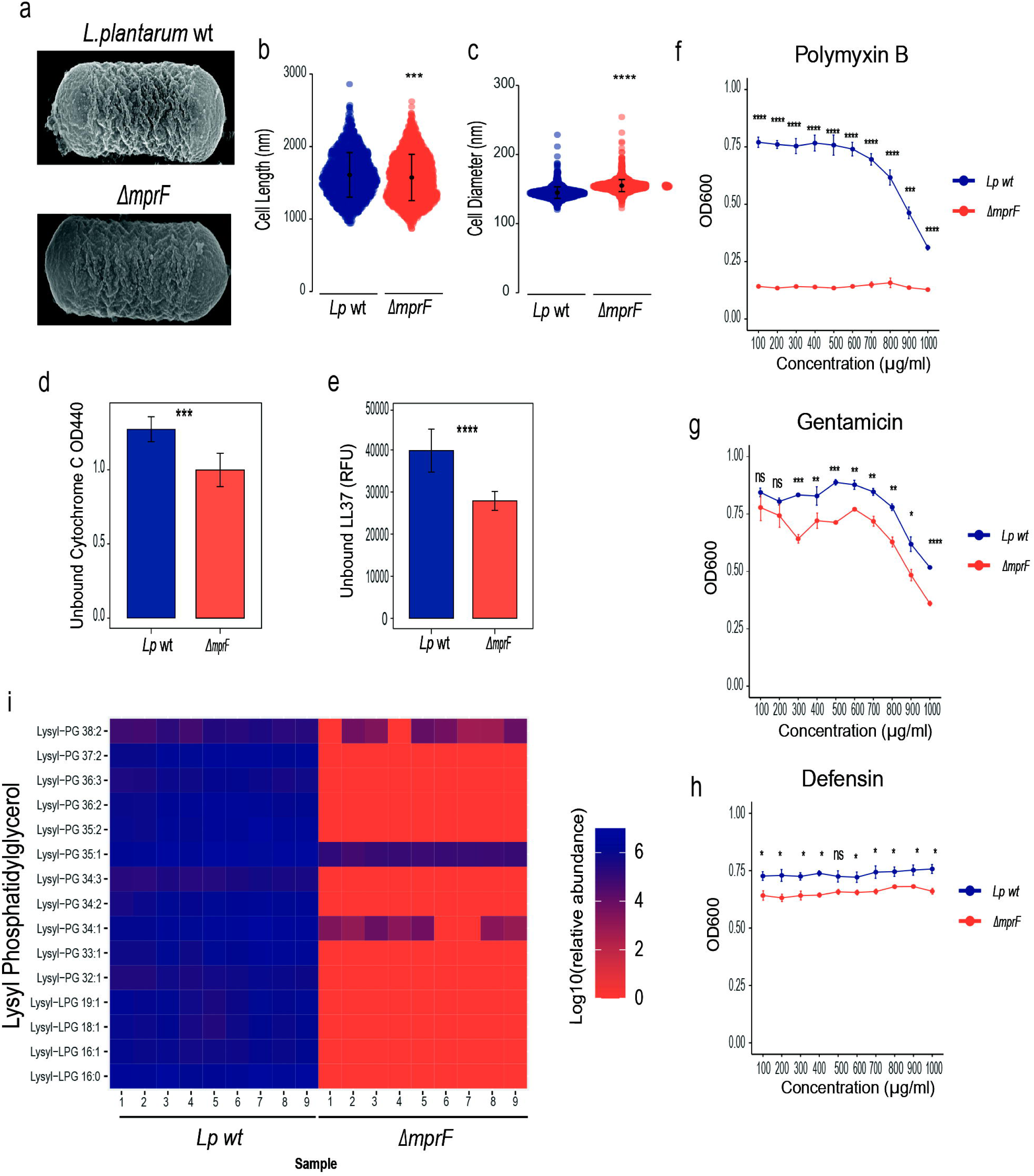
*L. plantarum* MprF mediates lipid lysylation and resistance to AMPs. (**a**) Scanning electron microscopy images of *L. plantarum* wild-type and Δ*mprF* mutant. (**b**) Cell length and (**c**) cell diameter of *L. plantarum* wild-type (n= 1372) and *L. plantarum* Δ*mprF* mutant (n=1927). Individual dots show single cell record. Violin dot plots show median and interquartile ranges. **(d, e)** Binding of *L. plantarum* wild-type and *L. plantarum* Δ*mprF* mutant cells to Cytochrome C (**d**) and to fluorescently labeled antimicrobial peptide LL37 (**e**) (n=3 independent experiments). Data show quantity of remaining cytochrome C (quantified by measuring OD440) or fluorescently labelled antimicrobial peptide LL37 (quantified by measuring fluorescence and expressed as Relative Fluorescent Units, (RFU) in the solution after incubation with indicated bacteria. Bar plots show mean and SEM. (**f-h**) Antibiotic Inhibitory Assay (AIA) of *L. plantarum* wild-type and *L. plantarum* Δ*mprF* mutant in MRS media supplemented with Polymyxin B (**g**), Gentamicin (**h**), and Defensin (**i**) (n = 3 independent experiments). (**i**) Heat map showing quantification of lysylated phospholipids in *L. plantarum* wild-type and *L. plantarum* Δ*mprF* mutant by LC-MS (n=3 independent experiments with 3 replicates each). Blue color illustrates high abundance, red color – absence/low abundance or a particular lipid. *P < 0.05, **P < 0.01, ***P< 0.001, ****P < 0.0001. Kruskal-Wallis and Bonferroni post hoc tests were used for the statistical analysis.

Next, we asked which defense mechanisms are in control of the Δ*mprF* mutant. We tested melanisation and, as expected from previous studies [25], a melanisation-defective *PPO1*^Δ^*2*^Δ^ fly mutant was more sensitive to wild-type *S. aureus*; however, the Δ*mprF* mutant remained similarly attenuated in both *PPO1*^Δ^*2*^Δ^ and wild-type flies (Fig. S1b). We found a similar result with hemocyte-deficient flies (Fig. S1c), suggesting that melanisation and hemocytes are not involved in the control of the Δ*mprF* mutant. Next, we tested the role of Toll pathway in the control of Δ*mprF* mutant by infecting *PGRP-SA* mutant flies deficient for the peptidoglycan-recognition receptor. Interestingly, the *S. aureus* Δ*mprF* mutant was as virulent as wild-type *S. aureus* to *PGRP-SA* deficient flies as illustrated by survival (Fig. 1c) and within the host pathogen growth (Fig. 1b). Thus, the Toll pathway is important to control infection by the Δ*mprF* mutant, likely because of the Δ*mprF* mutant’s sensitivity to Toll pathway effectors. To find these effectors, we infected flies lacking AMPs and found they were significantly more susceptible to Δ*mprF* mutant as compared to wild-type flies (Fig. 1d). Also, AMP-deficient flies in contrast to wild-type flies were not able to control the proliferation of Δ*mprF* mutant (Fig. 1e). Thus, AMPs at least in part are responsible for the control of the Δ*mprF* mutant. Additionally, we tested the effect of *mprF* mutation on the Toll pathway activation by monitoring the expression of *Drosomycin,* an AMP controlled by the Toll pathway, in infected flies. We found that infection of wild-type flies with the *S. aureus* Δ*mprF* mutant triggered significantly higher *Drosomycin* expression as compared to the infection with wild-type *S. aureus* (Fig. 1f). Such differences in *Drosomycin* expression were not observed in the *PGRP-SA* mutant, suggesting that they are caused by the differential Toll pathway activation by the two *S. aureus* strains (Fig. 1f). Thus, MprF mediates *S. aureus* virulence to *Drosophila* by protecting the pathogen from the effectors of Toll pathway and by reducing the activation of Toll pathway. Also, these results prove that lipid modifications by MprF are relevant for pathogen-fruit fly interactions motivating us to explore the function of MprF in *Drosophila* gut commensals.

### *L. plantarum mprF* mediates lipid lysylation and resistance to AMPs

Considering our recent findings that microbiota resistance to AMPs is necessary to maintain stable associations with the host particularly during immune challenge [14], we asked if *mprF* might be necessary for commensal resilience in the host gut. To address this question, we selected one of the prevalent *Drosophila* microbiota members – *L. plantarum* – as a model and used Cas9-based editing to generate *L. plantarum* Δ*mprF* mutant with the entire *mprF* open reading frame deleted (Fig. S2a-S2d). Due to good genetic tractability, we used the *L. plantarum* WCFS1 strain (also called NCIMB 8826). *L. plantarum* Δ*mprF* mutant did not show any growth differences compared to wild-type in MRS medium (Fig. S3). Also, we did not see any obvious morphological alterations with SEM (Fig. 2a). However, on average the cell length of *L. plantarum* Δ*mprF* mutant was reduced, while cell width increased (Fig. 2b-c). Given that in several bacteria mutations in *mprF* were shown to alter surface charge and increase binding of cationic AMPs to bacteria, we tested whether this is the case for the *L. plantarum* Δ*mprF* mutant. We measured the amount of cationic cytochrome C and fluorescently-labeled cationic AMP 5-FAM-LC-LL37 that remained in the solution after incubation with wild-type and *L. plantarum* Δ*mprF* cells. We detected cytochrome C and LL37 (Fig. 2d-e) in significantly lower amounts in the supernatants of *L. plantarum* Δ*mprF* as compared to wild-type *L. plantarum*, indicating an increased binding and negative cell surface charge in *L. plantarum* Δ*mprF*. Consistent with increased negative cell surface charge, the *L. plantarum* Δ*mprF* was more sensitive than wild-type *L. plantarum* to cationic antimicrobial peptide polymyxin B (Fig. 2f), antibiotic gentamicin (Fig. 2g), and insect AMP defensin (Fig. 2h) across a range of concentrations. We confirmed the increased sensitivity of *L. plantarum* Δ*mprF* to gentamycin by measuring the growth of bacteria in the presence of a specific concentration of antibiotic (Fig. S3a). Additionally, with this assay we detected increased sensitivity also to daptomycin (Fig. S3b) and nisin (Fig. S3c). Given that in other bacteria MprF neutralizes the membrane surface charge and provides AMP resistance by catalyzing the modification of the negatively charged lipid phosphatidylglycerol (PG) with L-lysine, we reasoned that *L. plantarum* MprF has a similar function. If this is true, we should expect reduced abundance of lysylated lipids in *L. plantarum* Δ*mprF*. Our lipidomic analysis indeed identified several Lys-PG species in wild-type *L. plantarum*. Most of them, however, were not detected in the *L. plantarum* Δ*mprF* strain, and those that were detected (Lys-PG 38:2, Lys-PG 35:1, Lys-PG 34:1) had significantly reduced abundance compared to wild-type *L. plantarum* (Fig. 2i). Thus, MprF is necessary for production of Lys-PG in *L. plantarum*, which reduces negative cell surface charge, binding of CAMPs, and increases resistance to cationic antimicrobials.

### MprF mediates *L. plantarum* persistence in the *Drosophila* gut

To test the in vivo importance of *mprF* for *L. plantarum*, we measured bacterial persistence in the gut during immune challenge (Fig. 3a). First, we exposed flies monocolonized with wild-type *L. plantarum* or the Δ*mprF* mutant to infection with the natural *Drosophila* gut pathogen *Pectobacterium carotovorum (Ecc15)*. While Δ*mprF* mutant and wild-type counts were similar in uninfected flies, we observed significantly reduced Δ*mprF* mutant counts 6 h and 24 h after infection in wild-type flies (Fig. 3b). However, in *Relish* or Δ*AMP* mutant flies, Δ*mprF* mutant loads were not significantly different from wild-type after infection at both time points tested (Fig. 3b), suggesting that AMPs regulated by the Imd pathway affect Δ*mprF* mutant abundance during intestinal inflammation. Moreover, we performed priming experiment (Fig. 3c), where *L. plantarum* wild-type and Δ*mprF* were introduced to the gut after infection. Again, by scoring bacterial abundance, we found that the Δ*mprF* mutant was as efficient as wild-type *L. plantarum* in gut colonization of flies that were not primed. In contrast, we detected significantly reduced ability of the mutant to colonize guts that were primed by infection at two time points tested (Fig. 3d). Importantly, the Δ*mprF* strain was able to colonize the guts of primed *Relish* or Δ*AMP* mutant flies to the same extend as wild-type *L. plantarum*, again pointing towards AMPs as regulators of Δ*mprF* mutant abundance. Additionally, we tested how genetic activation of the Imd pathway in the gut by *Imd* or *Relish* overexpression affects the persistence of the *L. plantarum* Δ*mprF* mutant in the gut. We found that while genetic activation of immune response in the gut had no effect on the abundance of wild-type *L. plantarum*, it significantly lowered the counts of Δ*mprF* mutant at 6, 24, and 48 h post colonization (Fig. 3e). We could restore the ability of the Δ*mprF* mutant to colonize guts of flies with genetically activated immune response by multi-copy plasmid-based expression of *mprF* in the Δ*mprF* mutant (Fig. 3f). These results together indicate that MprF-mediated resistance to host AMPs is necessary for *L. plantarum* persistence in the gut during intestinal inflammation.

**Figure 3.**
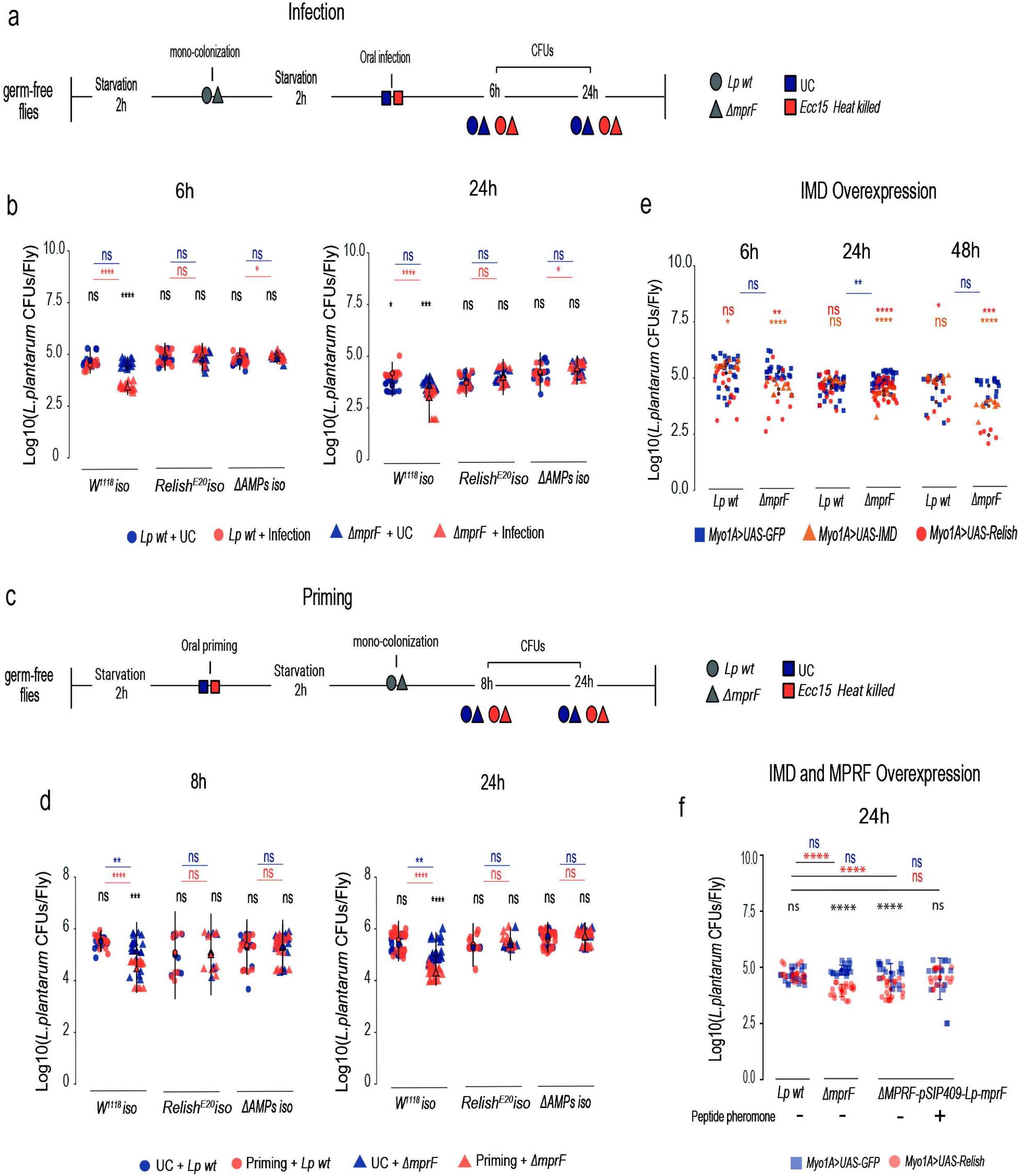
MprF mediates *L. plantarum* persistence in the *Drosophila* gut during inflammation. (**a**) Experimental design for monocolonization and infection protocol. (**b**) *L. plantarum wild-type* and Δ*mprF* loads in wild-type, *Relish^E20^*, and Δ*AMPs* flies at 6h and 24h after infection with *Ecc15* (n=14 independent samples per treatment with 5 flies per sample). (**c**) Experimental design for oral priming protocols (**d**) *L. plantarum wild-type* (n1) and Δ*mprF* (n2) loads in flies primed or not with *Ecc15* at 8h and 24h in wild-type, *Relish^E20^*, and Δ*AMPs* flies. Number of samples (n): 8h wild-type flies (n1=20, 20; n2=20, 16), *Relish^E20^* (n1=8, 8; n2=7, 8) Δ*AMPs* (n1=19, 22; n2=20, 20)., 24h wild-type (n1=20, 20; n2=20, 17) *Relish^E20^* (n1=6, 10; n2=7, 10), Δ*AMPs* (n1=21, 24; n2=21, 19). **(e)** *L. plantarum* wild-type (n1) and Δ*mprF* (n2) loads in *Myo1A-GAL4>UAS-GFP*, *Myo1A-GAL4>UAS-Relish,* and *Myo1A-GAL4>UAS-IMD* flies at 6h, 24h and 48h after colonization. Number of samples (n): *Myo1A-GAL4>UAS-GFP* (n1=26, 34, 14; n2= 22, 35, 14), *Myo1A-GAL4>UAS-Relish* (n1=12, 16, 5, n2=10, 33, 5) and *Myo1A-GAL4>UAS-IMD* (n1=9, 9, 10, n2=10, 10, 9), three time points are shown in the n sample. (**f**) Rescue of *L. plantatum* Δ*mprF* persistence by the overexpression of wild-type *mprF*. Loads of *L. plantarum* wild-type (n1), Δ*mprF* (n2), Δ*mprF-*pSIP409*-Lp-mprF* (n3), and Δ*mprF-*pSIP409*-Lp-mprF +* IP-673 peptide (n4). Number of samples (n): *Myo1A-GAL4>UAS-GFP* (n1=10, n2=10, n3=10, n4=8), *Myo1A-GAL4>UAS-Relish* (n1=10, n2=10, n3=10, n4=8). Individual dots show bacterial load per 5 female flies. Lines show median and interquartile ranges (IQR). *P < 0.05, **P < 0.01, ***P< 0.001, ****P < 0.0001. Kruskal-Wallis and Bonferroni post hoc tests were used for the statistical analysis.

### *L. plantarum* MprF confers Lys-PG synthesis and resistance to antibiotics in *E. coli*

In order to further characterize the function of *L. plantarum* MprF, we expressed the *L. plantarum mprF* gene in the heterologous host *E. coli*. *E. coli* lacks *mprF*-related genes and does not produce Lys-PG. Yet, *E. coli* contains PG, the putative Lys-PG precursor. The *L. plantarum mprF* was cloned in a multi-copy plasmid under an arabinose-inducible promoter. *E. coli* was transformed with the plasmid and *mprF* expression was induced with L-arabinose. A culture without arabinose was used as a control. Lipid analysis confirmed that *L. plantarum mprF* expression in *E. coli* leads to the synthesis of several Lys-PG species that are normally not produced by *E. coli* (Fig. 4a). Thus, *L. plantarum* MprF is necessary and sufficient for Lys-PG production in *E. coli*. Given that a prominent role of Lys-PG is to neutralize cell surface charge, we investigated whether increased synthesis of Lys-PG by *L. plantarum mprF* expression affects binding of cationic molecules to *E. coli*. We incubated *E. coli* expressing *mprF* and *E. coli* not expressing *mprF* with cytochrome C (Fig. 4b) or with fluorescently labelled LL37 peptide (Fig. 4c) and measured the amounts of both molecules that remained in the solution. We detected cytochrome C and LL37 (Fig. 4b-c) in significantly higher amounts in the supernatants of *E. coli* expressing *mprF* as compared to *E. coli* not expressing *mprF*, indicating that *mprF* expression reduces binding of cationic molecules to *E. coli* cells. Consistent with this, *L. plantarum mprF* expression increased *E. coli* resistance to several antibiotics and AMPs, like polymyxin B, gentamicin, and cecropin (Fig. 4d).

**Figure 4.**
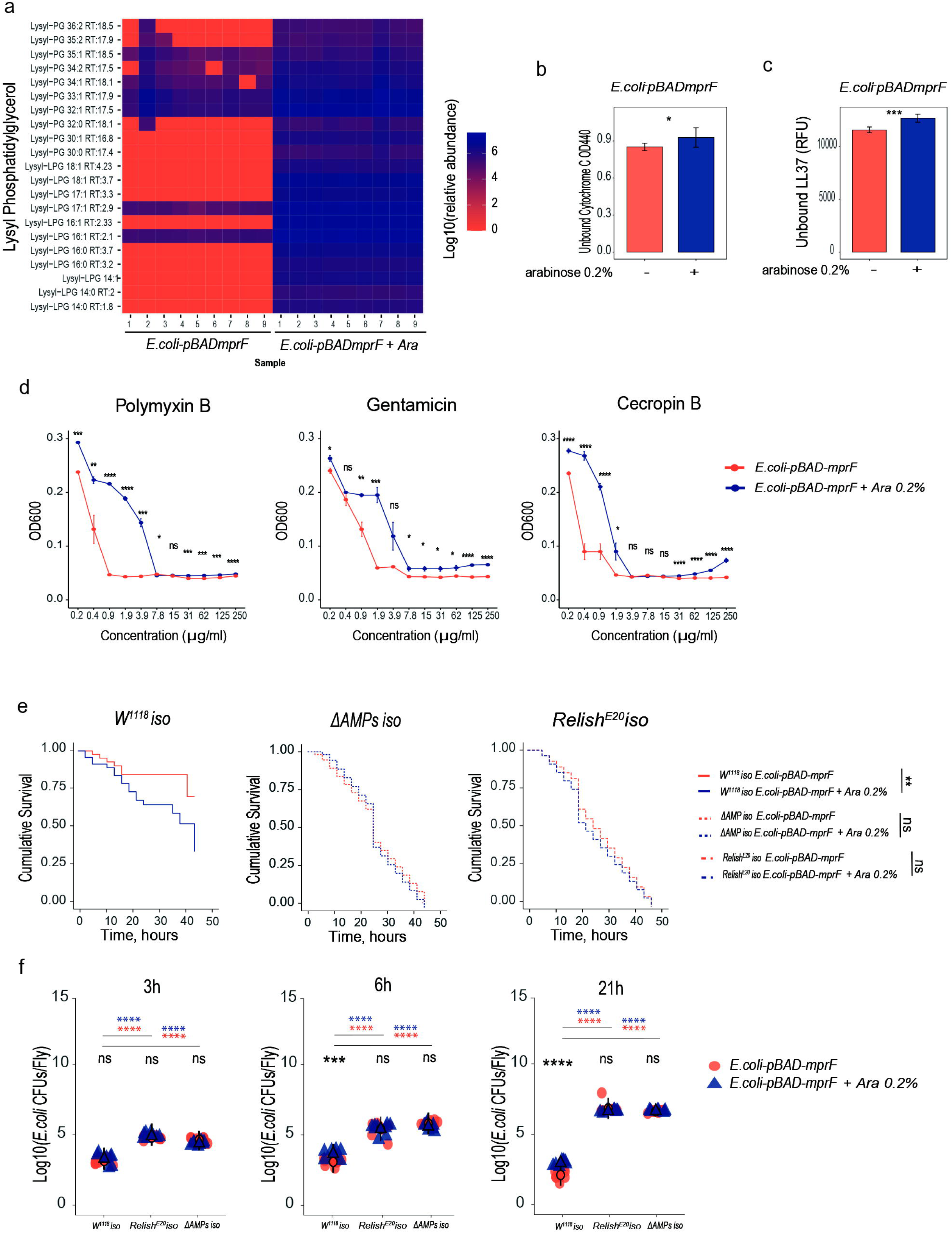
*L. plantarum mprF* confers Lys-lipid synthesis and antibiotic resistance in *E. coli*. (**a**) Heat map showing quantification of lysylated phospholipids in *E. coli* expressing (*E. coli-pBAD-mprF*+Ara) and not expressing *Lp*MprF (*E. coli-pBAD-mprF)*. Blue color illustrates high abundance, red color – absence/low abundance or a particular lipid. (**b, c**) Binding of *E. coli* cells expressing and not expressing *Lp*MprF to Cytochrome C (**b**) and to fluorescently labeled antimicrobial peptide LL37 (**c**) (n=3 independent experiments). Data show quantity of remaining cytochrome C (quantified by measuring OD440) or fluorescently labeled antimicrobial peptide LL37 (quantified by measuring fluorescence and expressed as Relative Fluorescent Units, RFU) in the solution after incubation with indicated bacteria. Bar plots show mean and SEM. (**d**) Antibiotics Inhibition Assay (AIA) on *E. coli* cells expressing and not expressing *Lp*MprF in LB media supplemented with Polymyxin B, Gentamicin, and Cecropin B (n = 3 independent experiments). (**e**) Survival rates of *w^1118^ iso,* Δ*AMP* and *Relish^E20^* flies infected with *E. coli* expressing (*E. coli-pBAD-mprF*+Ara) and not expressing *Lp*MprF (*E. coli-pBAD-mprF).* (**f**) Loads of *E. coli* expressing (*E. coli-pBAD-mprF*+Ara, n2) and not expressing *Lp*MprF (*E. coli-pBAD-mprF, n1)* in *w^1118^ iso* (n1=9, 9, 9, n2=8, 8, 9), *Relish^E20^* (n1=9, 9, 9, n2=9, 9, 9), and Δ*AMP* flies (n1=9, 9, 9, n2=9, 9, 9). Three time points per bacteria are shown in the n sample. *Lp*MprF expression in *E. coli-pBAD-mprF* was induced with L-arabinose 0.2%. In dot plots, median and interquartile ranges are shown (IQR); whiskers show either lower or upper quartiles or ranges. Mean and SEM are shown. *P < 0.05, **P < 0.01, ***P< 0.001, ****P < 0.0001. Kruskal-Wallis and Bonferroni post hoc tests were used for the statistical analysis.

Additionally, we investigated whether *L. plantarum* MprF can protect *E. coli* from AMPs in vivo. We performed systemic infections of wild-type, *Relish* and Δ*AMP* mutant flies with control *E. coli* and with *E. coli* expressing *L. plantarum mprF*. While control *E. coli* had little effect on survival of wild-type flies, *mprF* expression significantly increased *E. coli* virulence as illustrated by the increased proportion of dead flies (Fig 4e). Survival differences caused by infections with the two *E. coli* strains were not detected in Δ*AMP* or *Relish*-deficient flies both of which showed increased susceptibility to infections with the two *E. coli* strains (Fig. 4e). Consistent with survival results, we detected significantly more CFUs of *E. coli* expressing *mprF* relative to control *E. coli* at 6 h and 21 h post infection in wild-type flies (Fig. 4f). While *E. coli* load was significantly higher in Δ*AMP* and *Relish*-deficient flies, there was no difference in the amount of control and *mprF*-expressing *E. coli*. These results suggest that *L. plantarum mprF* expression confers increased virulence to *E. coli* only in the presence of Imd-regulated AMPs, likely by increasing *E. coli* resistance to these AMPs.

### MprF affects bacterial immunostimulatory properties by limiting the release of PGN fragments

The fact that infection with an *S. aureus* Δ*mprF* mutant as compared to wild-type *S. aureus* resulted in elevated Toll pathway activation motivated us to test the immunomodulatory properties of the *L. plantarum* Δ*mprF* mutant. For this purpose, we assessed Imd pathway activation by measuring the expression of Imd pathway-regulated AMP *Diptericin (Dpt)* in the guts of flies colonized with either wild-type *L. plantarum* or the Δ*mprF* mutant. We found that flies monocolonized with the *L. plantarum* Δ*mprF* mutant showed significantly higher *Dpt* expression in the guts compared to flies colonized with wild-type *L. plantarum*. This response is dependent on the activation of the Imd pathway, as it is abolished in the *Relish* mutant (Fig. 5a). This finding demonstrates a dual role of MprF in *L. plantarum*: first, it modifies the bacterial cell surface, thereby facilitating resistance to cationic antimicrobials; second, it reduces bacterial sensing by the Imd pathway, thus mediating evasion of the immune response.

**Figure 5.**
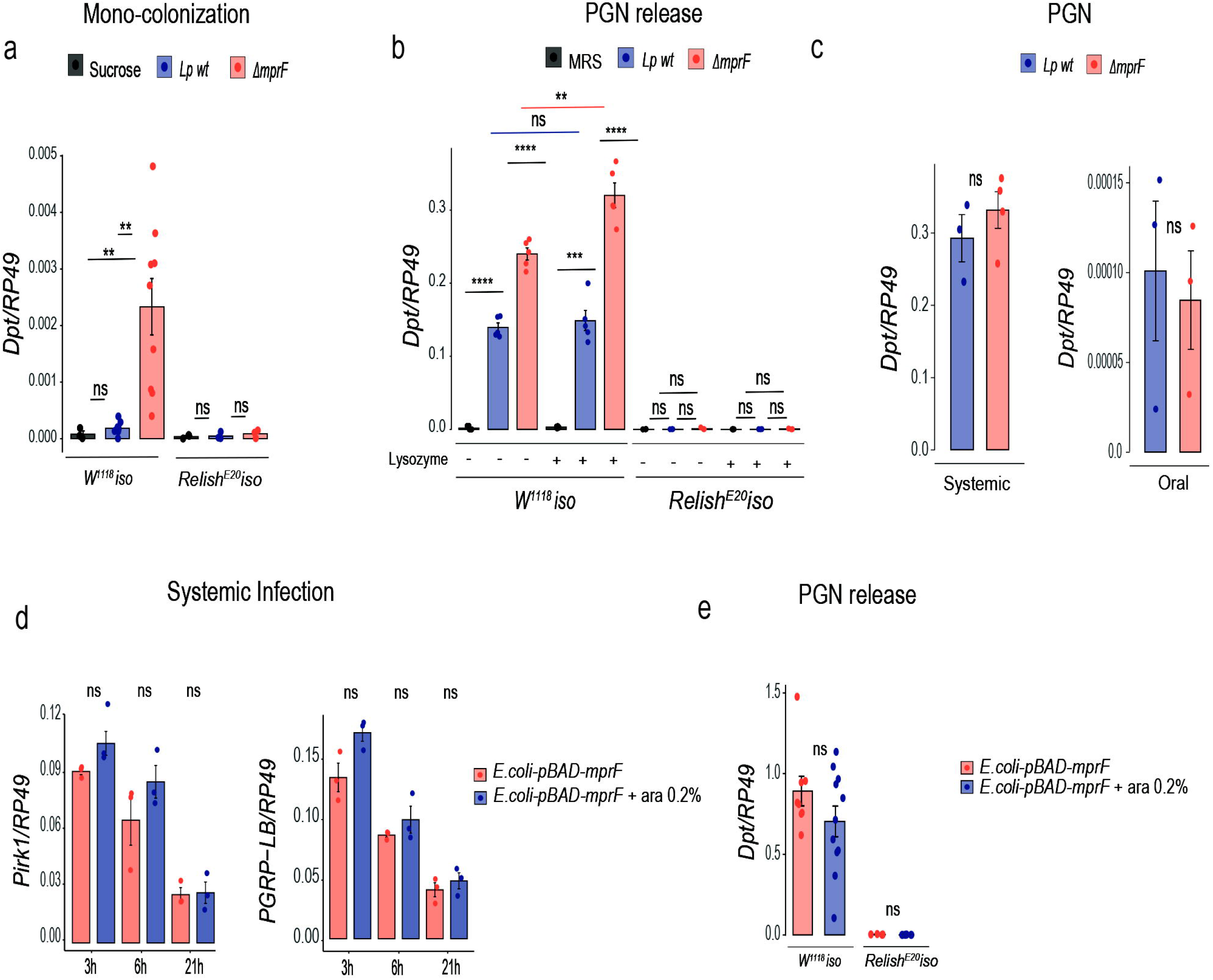
MprF affects bacterial immunostimulatory properties by limiting the release of PGN fragments. (**a**) Intestinal *Diptericin A* gene expression 5 days post colonization with *L. plantarum wild-type* (n1) *and* Δ*mprF mutant* (n2) or treatment with sucrose 2.5% only (n3) in wild-type (n1=9, n2=9, n3=3) and *Relish^E20^*flies (n1=3, n2=3, n3=3). (**b**) Systemic *DptA* gene expression 4h after injection of supernatants from *L. plantarum wild-type* (n1), Δ*mprF mutant* (n2) or MRS (n3) without chicken egg lysozyme treatment in wild-type (n1=5, n2=5, n3=5) and *Relish^E20^* flies (n1=3, n2=3, n3=3), or after treatment with chicken egg lysozyme, in wild-type (n1=5, n2=5, n3=5) and *Relish^E20^* flies (n1=3, n2=3, n3=3). (**c**) Systemic *DptA* gene expression 4h after injection (n1=3, n2=4) and intestinal *DptA* expression 4h after ingestion (n1=3, n2=3) of purified PGN from *L. plantarum* wild-type (n1) *or* Δ*mprF mutant* (n2). (**d**) Systemic *Pirk* and *PGRP-LB* gene expression after systemic infection with *E. coli* expressing (*E. coli-pBAD-mprF*+Ara, n2) and not expressing *Lp*MprF (*E. coli-pBAD-mprF, n1)* at 3h (n1=3, n2=3), 6h (n1=3, n2=3) and 21h (n1=3, n2=3) post infection. (**e**) Systemic *DptA* gene expression 4h after injection of supernatants from *E. coli* expressing (*E. coli-pBAD-mprF*+Ara, n2) and not expressing *Lp*MprF (*E. coli-pBAD-mprF, n1)* in wild-type (n1=8, n2=11) and *Relish^E20^*flies (n1=4, n2=4). *Lp*MprF expression in *E. coli-pBAD-mprF* was induced with 0.2% L-arabinose. Individual dots show gene expression per 20 female guts (intestinal expression) or 5 whole female flies (systemic expression). Bar plots show mean and SEM. *P < 0.05, **P < 0.01, ***P< 0.001, ****P < 0.0001. Kruskal-Wallis and Bonferroni post hoc tests were used for the statistical analysis.

Given that PGN fragments are the major elicitors of the Imd pathway in flies, we hypothesized that the Δ*mprF* mutant, while being exposed to cationic AMPs, releases cell wall fragments which elicit strong Imd pathway activation. To test this possibility, we cultivated the wild-type and *L. plantarum* Δ*mprF* mutant in culture medium supplemented with or without lysozyme, a cationic antimicrobial. We collected cell-free culture supernatants of wild-type *L. plantarum* and the Δ*mprF* mutant, injected them into flies and measured *Dpt* expression to estimate Imd pathway activation. Injection of both supernatants resulted in *Dpt* expression, confirming that bacteria indeed discharge PGN fragments during growth (Fig. 5b). However, while lysozyme treatment had no significant effect on *Dpt* expression induced by the injection of supernatants from wild-type *L. plantarum*, the supernatant of the Δ*mprF* mutant treated with lysozyme triggered significantly stronger *Dpt* expression relative to the supernatant from an untreated culture (Fig. 5b). We did not detect any *Dpt* expression in the *Relish* mutant upon supernatant injection, confirming that the triggered response is Imd pathway-dependent. Notably, the supernatants of both lysozyme-treated and untreated Δ*mprF* mutant cultures elicited significantly higher *Dpt* expression as compared to that of wild-type *L. plantarum*, suggesting an increased release of immunostimulatory PGN fragments by the Δ*mprF* mutant.

Next, we asked whether potential structural differences between *L. plantarum* wild-type and Δ*mprF* mutant PGN could also contribute to the variation in Imd-elicited responses. Therefore, we purified cell wall fractions from the wild-type *L. plantarum* and Δ*mprF* mutant bacteria and compared *Dpt* expression upon their injection and feeding in flies. Injection of equal amounts of purified cell wall fractions from both strains triggered comparable level of Imd pathway activation (Fig. 5c). A similar result was observed with intestinal Imd pathway activation induced by feeding flies with purified cell wall fractions (Fig. 5c). Altogether, these data confirm that differences in the Imd-triggered response elicited by wild-type and Δ*mprF* mutant can be linked to varying doses of discharged PGN fragments rather than to structural differences in PGN.

Given our findings that *mprF* expression promotes lipid lysylation and increases *E. coli* resistance to AMPS, we tested whether *L. plantarum mprF* expression will also affect immunomodulatory properties of *E. coli*. As shown in Fig. 5d, expression of two Imd-pathway regulated genes, *Pirk* and *PGRP-LB*, was not significantly different between flies systemically infected with control *E. coli* and *E. coli* expressing *mprF*. Similarly, overexpresion of *mprF* in *E. coli* did not significantly affect the release of PGN fragments, as injection of supernatant from *mprF*-expressing *E. coli* resulted in a similar level of Imd pathway activation as observed with the injection of control supernatant (Fig. 5e). Hence, the immunomodulatory properties of *E. coli* in contrast to AMP susceptibility were not significantly altered by *mprF* overexpression.

### LTA production is altered in the *L. plantarum* Δ*mprF* mutant

Considering recent finding that MprF affects the length of lipoteichoic acids (LTAs) in *B. subtilis* and *S. aureus* [55], we tested whether MprF has a similar role in *L. plantarum*. To this end, we compared LTA profiles of wild-type *L. plantarum* and Δ*mprF* mutant using crude bacterial extracts and a western blot with a monoclonal antibody against Gram-positive LTA. As shown in Fig. 6a, wild-type *L. plantarum* and the Δ*mprF* mutant have distinct LTA profiles, where LTA from Δ*mprF* mutant migrated faster, indicating reduced size. To confirm that our western blot indeed detects differences in LTA profiles and not in any other components present in the bacteria extracts, we performed western blot with purified LTA and obtained similar results (Fig. S4). Additionally, we applied purified *L. plantarum* LTA after de-O-acylation by hydrazine-treatment [61] to a Tris-tricine-PAGE analysis (Fig. 6b; full length gel shown in Fig. S5) specifically optimized for TA analysis [62, 63]. We could confirm the reduced overall length of LTA in the Δ*mprF* mutant with this method, too. ^1^H NMR spectra recorded from both native (Fig. 6c) and hydrazine-treated LTA (Fig. 6d) of the two strains showed that the overall structural composition is not altered between the wild-type and the mutant *L. plantarum* strain. Therefore, the reduced size of LTA in the Δ*mprF* mutant is not due to general structural changes but is likely because of the reduced size of the polymeric LTA chain.

**Figure 6.**
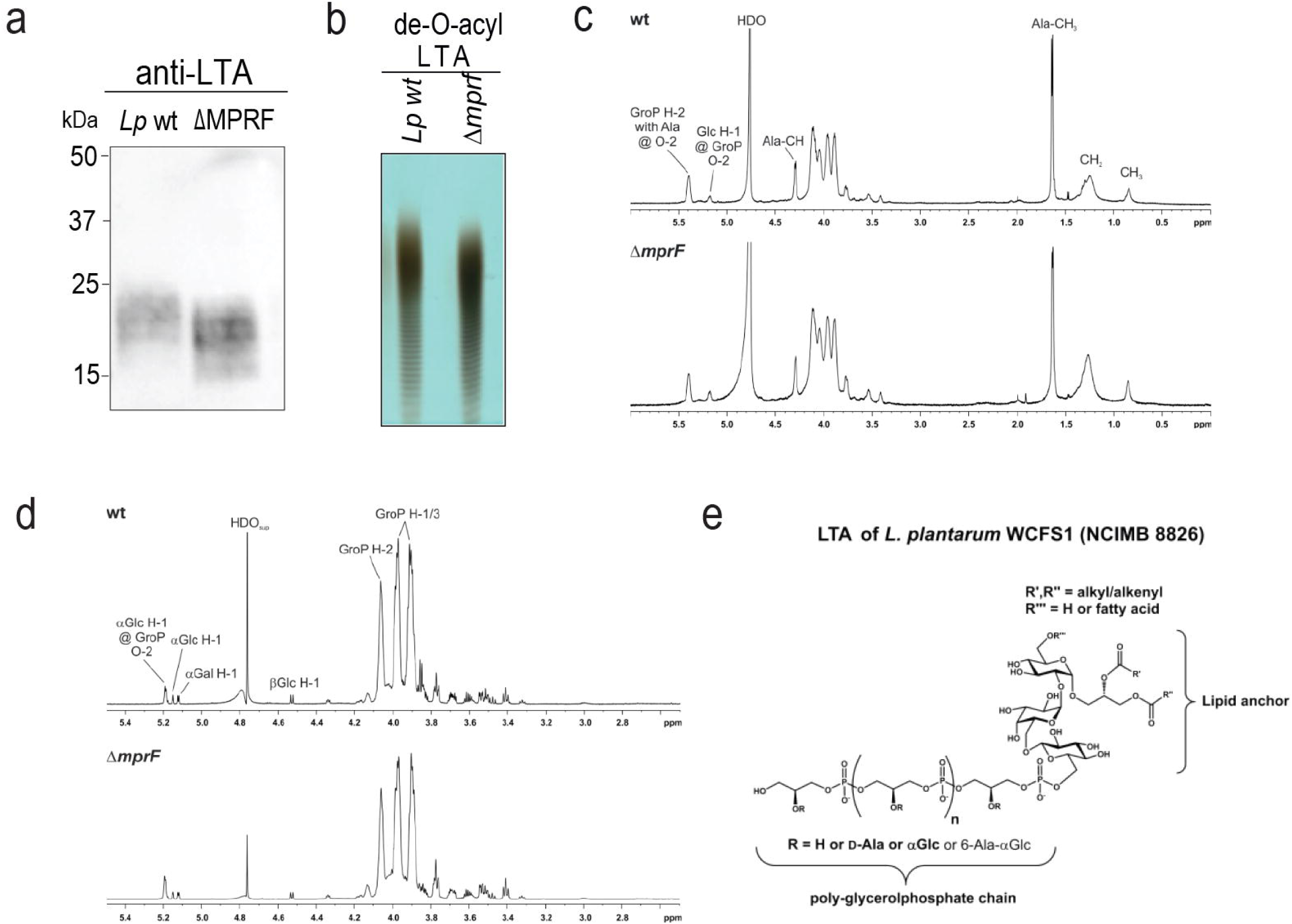
LTA chain length is partially reduced in the *L. plantarum* Δ*mprF* mutant. (**a**) LTA profile of *L. plantarum* wild-type and Δ*mprF* mutant detected with Western blot and anti-LTA MAb at a 1:1,000 dilution. (**b**) Profile of de-O-acyl LTA visualized by Tris-tricine-PAGE with combined alcian blue and silver staining (full length gel is shown in Fig. S5). (**c**, **d**) ^1^H NMR analysis of native (c) and de-O-acyl (d) LTA. Shown are ^1^H NMR spectra (δ_H_ 6.0–0.0 ppm (native) or δ_H_ 5.5–2.5 ppm (de-O-acyl)) recorded in D_2_O at 300 K. (**e**) Chemical structure of LTA isolated from *L. plantarum* WCFS1 (NCIMB 8826). Position of the third fatty acid (R’’’) was adopted from ref. [66].

Since the structure of LTA from *L. plantarum* strain WCFS1 (NCIMB 8826) has not been described yet, we performed a full NMR analysis. Analyzing the de-O-acylated LTA after hydrazine treatment enables a much better resolution of the carbohydrate parts of the molecule, especially the ones of the glycolipid anchor. These sugars are almost undetectable in NMR spectra of native LTA due to the known formation of micelles when LTA is dissolved in aqueous solutions [61, 64]. The structural model for *L. plantarum* strain WCFS1 (NCIMB 8826) LTA is depicted in Fig. 6e, NMR chemical shift data for the hydrazine-treated LTA are listed in Table S1. The observed LTA structure is in line with described structures or structural features of other *L. plantarum* strains. The glycolipid anchor consists of the trisaccharide βGlc*_p_*-(1→6)-αGal*_p_*-(1→2)-αGlc*_p_* that is 1,3-linked to a diacylglycerol as it has been described for LTA from *L. plantarum* strain K8 (KCTC 10887BP) [65]. The presence of this glycolipid in either di- or tri-acylated form has also been reported for *L. plantarum* strain IRL-560 [66]. In this study, we could unequivocally proof by an ^1^H,^31^P-HMQC experiment (Fig. S6) that the poly-glycerolphosphate chain is coupled to the O-6 position of the βGlc*_p_*residue. As described for strain K8 [65] and NC8 [67] the major substituents at the O-2 position of the glycerolphosphate moities are alanine (Ala) and αGlc*_p_* residues. In addition, we have evidence for a small proportion of 6-Ala-αGlc*_p_* as additional substituent (Fig. S7) like it has been described recently for strain NC8 [67]. However, we found no evidence for a putative αGal*_p_*-substitution as mentioned for strain K8 [65]. In conclusion, we describe the LTA structure of *L. plantarum* type strain WCFS1 and show that *mprF* deficiency doesn’t alter the overall structural composition of LTA but likely affects the length of the polymeric LTA chain.

## Discussion

As a starting point for this work, we used the existing *mprF* mutants to test the relevance of *mprF* in a *Drosophila* model and investigated whether MprF is required for pathogen virulence in a *Drosophila* model and whether this is linked to immune evasion. Using *S. aureus* as a pathogen efficiently infecting fruit flies, we found that MprF is required for the virulence of this pathogen, as flies infected with an *S. aureus* Δ*mprF* mutant survived significantly longer compared to counterparts infected with wild-type *S. aureus*. The virulence of the Δ*mprF* mutant was restored in flies not able to sense Gram-positive pathogens and initiate Toll-dependent AMP response, suggesting that AMPs induced by the Toll pathway likely clear Δ*mprF* mutant. In line with this, the Δ*mprF* mutant was more virulent to AMP-deficient flies than to wild-type flies. However, Δ*AMP* mutants were not as sensitive as *PGRP-SA*-deficient flies to an *S. aureus* Δ*mprF* mutant, suggesting that while AMPs control the Δ*mprF* mutant, there are additional Toll pathway-regulated effectors involved. Since we ruled out the melanisation response in the control of the Δ*mprF* mutant, peptides from the Bomanin family, as major Toll pathway effectors [68], are attractive candidates that could be tested. Overall, our results support a role of MprF in *S. aureus* virulence to *Drosophila* by facilitating pathogen evasion of Toll pathway-dependent effectors.

Our results with existing *S. aureus* Δ*mprF* mutant identified MprF as a relevant factor in pathogen-*Drosophila* interactions. Being motivated by these findings, we investigated the role of MprF in commensal-host interactions, using the prominent fruit fly gut microbe *L. plantarum* as a model. Phenotypic characterization of the *L. plantarum* Δ*mprF* mutant that we generated revealed similarity to *mprF* mutants in other bacteria. Namely, the *L. plantarum* Δ*mprF* mutant exhibited deficiency in the synthesis of Lys-PG, increased negative cell surface charge, increased binding of cationic molecules, and enhanced sensitivity to AMPs. Our in vivo analysis demonstrated that the abundance of the *L. plantarum* Δ*mprF* mutant significantly declined in fly guts during infection- or genetically-induced immune response. Such decline is not due to general inability of the mutant to colonize the gut but is attributed to mutant’s sensitivity to host AMPs, since the abundance of the Δ*mprF* mutant was not affected by infection in AMP-deficient flies and Δ*mprF* mutant colonized the guts of uninfected flies as efficiently as wild-type *L. plantarum*. Overall, we demonstrated that MprF besides its well-described role in pathogen resistance to AMPs and virulence is also an important factor mediating commensal resilience during inflammation via AMP resistance. Thus, our work further advances our understanding of how host-microbiota homeostasis is maintained during infection-induced inflammation.

Importantly, the other recent studies further support the role of MprF in microbiota persistence in the gut. For example, a metagenome-wide association (MGWA) study identified multiple bacterial genes, including *mprF*, that are significantly correlated with the level of colonization [69]. Subsequent analyses confirmed that an *mprF* transposon insertion mutant of *Acetobacter fabarum* showed decreased persistence within the flies [69]. However, it has not been tested whether this phenotype is due to mutant’s sensitivity to host AMPs. Another study compared the evolutionary trajectory of *L. plantarum* in the fly food and inside the flies. They showed that *L. plantarum* populations that evolved in the presence of fruit flies were repeatedly affected by non-synonymous mutations in the *mprF* gene [70]. Similarly, mutations in *mprF* gene were identified in *E. faecalis* during experimental evolution via serial passage in *Drosophila* [71]. These studies suggest that *mprF* is under selection in the host environment. The significance of these mutations for bacterial association with flies, however, has not been tested. *E. coli* strain that is made resistant to AMPs by *mcr-1* gene expression similarly showed better ability to persist in the mouse gut, highlighting an important role of AMP resistance for commensal lifestyle [72].

We noticed that flies colonized with the *L. plantarum* Δ*mprF* mutant exhibited elevated gut AMP response, indicating that besides well-characterized role in AMP resistance, MprF has previously undescribed function in modulating bacterial immunostimulatory properties. Hence, MprF confers two relevant for commensal-host association functions: AMP resistance and immune evasion. Whether MprF mediates immune evasion similar to AMP resistance mechanism via production of Lys-PG remains to be tested. However, our experiments with heterologous expression of MprF in *E. coli* point towards Lys-PG independent role of MprF in immunomodulation. Specifically, the facts that MprF induces Lys-PG synthesis in *E. coli* and increases resistance to AMPs but doesn’t affect immunostimulatory properties support a possibility that immunomodulatory properties are not linked to MprF-mediated PG lysylation.

Being motivated by recent studies on MprF’s role in LTA production in *S. aureus* and *B. subtilis* [55], we analysed the LTA profile in the *L. plantarum* Δ*mprF* mutant. Consistent with the published observation in *S. aureus*, we similarly detected to some extent reduced size of LTA in the *L. plantarum* Δ*mprF* mutant as compared to wild-type *L. plantarum*. However, since the overall structure of *L. plantarum* WCFS1 (NCIMB 8826) LTA – which was found to be very similar to LTAs described for other *L. plantarum* strains – was not altered, it is rather unlikely that this LTA size reduction significantly alters the physiology of *L. plantarum*. Since *E. coli* lacks LTA, MprF presence in *E. coli* would not change LTA but could increase Lys-PG synthesis, thus providing a potential explanation as to why MprF overexpression affects *E. coli* AMP resistance but not immunostimulatory properties. Furthermore, D-alanylation of TAs was shown to prevent the discharge of immunostimulatory PGN fragments by *L. plantarum* and activation of fly immune response [44]. A similar phenotype that we described here for the *L. plantarum* Δ*mprF* mutant, suggests a potential link between reduced LTA size and enhanced release of PGN fragments. Additionally, LTA might affect the PGN’s accessibility to recognition by PRRs, as was reported for WTAs [73]. Alternatively, recent finding that D-Ala-LTAs act as direct bacterial cues for *Drosophila* larvae to initiate growth-promoting effect [67], raises a possibility that LTAs instead of affecting PGN availability/accessibility might be direct signals sensed by *Drosophila* intestinal epithelial cells. It is also possible that MprF’s effect on resistance to AMPs and antibiotics is not exclusively mediated by Lys-PG synthesis but rather by modifications of LTAs. This seems to be the case for daptomycin, for example [55]. Yet, both functional consequences and the mechanisms of MprF’s contribution to LTA synthesis require further investigation.

There is accumulating evidence that factors originally implicated in pathogen immune evasion and virulence are also essential for commensal persistence within the host. Besides MprF described here, we and others previously illustrated the role of the *dlt* operon in *L. plantarum* resilience during inflammation [14, 44]. Similarly, LPS-mediated resistance to AMPs was identified as one of the major virulence factors of *Providencia alcalifaciens* in *Drosophila* [74] and as an essential mechanism of AMP resistance and gut colonization by the insect symbiont *Caballeronia insecticola* [75]. These studies further support the notion that both host-symbiont and host-pathogen associations are governed by a shared molecular dialogue [76], which we are just beginning to understand. This notion has important practical implications for the development of antivirulence strategies targeting pathogen immune evasion factors [77]. Considering an essential role of some of these factors for microbiota persistence within the host, antivirulence approaches targeting pathogen evasion factors should consider potential impact of such treatments on host microbiota and its stability.

## Materials and Methods

### *Drosophila* stocks and rearing

The following *Drosophila* stocks used in this study were kindly provided by Dr. Bruno Lemaitre: DrosDel *w^1118^ iso*; *Canton S*; *Oregon R*; *yw*; *PGRP-SA^Seml^*; *PPO1*^Δ^*2*^Δ^ *iso*; *hml-Gal4*; *UAS-bax*; *Relish^E20^ iso*; Δ*AMP* iso; *UAS-Relish*; *UAS-Imd*; *w; Myo1A-Gal4, tubGal80TS, UAS-GFP*. Flies stocks were routinely maintained at 25 °C, with 12/12 hours dark/night cycles on a standard cornmeal-agar medium: 3.72 g agar, 35.28 g cornmeal, 35.28 g inactivated dried yeast, 16 mL of a 10% solution of methylparaben in 85% ethanol, 36 mL fruit juice, and 2.9 mL 99% propionic acid for 600 mL. Food for germ-free flies was supplemented with ampicillin (50 μg/mL), kanamycin (50 μg /mL), tetracyclin (10 μg/mL), and erythromycin (10 μg /mL). Fresh food was prepared weekly to avoid desiccation.

### Bacterial strains and survival experiments

Bacterial strains used in this study and their growth conditions are listed in Table S2. Systemic infections (septic injury) were performed by pricking adult flies (5 d to 10 d old) in the thorax with a thin needle previously dipped into a concentrated pellet of a bacterial culture. For infection assay, bacteria were pelleted by centrifugation and diluted with PBS to the desired optical densities at 600 nm (OD_600_). *S. aureus* was used at OD_600_=5, *E. coli* at OD_600_=300. In case of *S. aureus* infection, infected flies were kept at 25 °C overnight and switched to 29 °C for the rest of experiment. *E. coli*-infected flies were kept constantly at 29 °C. At least two vials of 20 flies were used for survival experiments, and survivals were repeated at least three times. Infected flies were maintained in vials with food without live yeasts during survival assays and until collection for bacterial load estimation or RNA extraction. In survival curves the cumulative data of three independent experiments are displayed.

### Generation of *L. plantarum* Δ*mprF* mutant

#### Plasmid construction

The primers/oligos used for cloning and the constructed plasmids are listed in Supplementary Tables S3 and S4. Genome editing in *L. plantarum* WCFS1 was performed using two *E. coli-Lactiplantibacillus* shuttle vectors. The first shuttle vector, pAA009 encodes SpyCas9, a tracrRNA and repeat-spacer-repeat array with a 30-nt spacer targeting the multiple peptide resistance factor (*mprF*) gene. The targeting spacer was added by restriction digestion of the backbone plasmid, pCB578, with PvuI-HF (NEB Cat. No. R3150S) and NotI-HF (NEB Cat. No. R3189S), followed by ligation (NEB Cat. No. M0370L) of the digested backbone with phosphorylated (NEB Cat. No. M0201L) annealed oligos oAA027-028. The second shuttle vector, pAA032, was used as the plasmid carrying a recombineering template to generate a clean deletion of the *mprF* gene in the WCFS1 strain. First, pCB591 was amplified with primers oAA094-099 to get a backbone fragment for pAA032, and *mprF* along with 250-bp homology arms flanking the start and stop codons was amplified with primers oAA097-098 using genomic DNA from WCFS1 as template. The PCR fragments were joined together using Gibson assembly kit (NEB Cat. No. E2611L) following the manufacturer’s instructions. Then, primers oAA033-034 were used to remove *mprF* using the Q5 site-directed mutagenesis kit (NEB Cat. No. E0554S) following the manufacturer’s instructions, yielding the final recombineering template pAA032. DH5α competent *E. coli* cells were used for both cloning steps and primers oAA038-039 were used for screening clones by colony PCR. Correct clones were confirmed by Sanger sequencing (Microsynth GmbH) and whole plasmid sequencing (Plasmidsaurus). After successful clones were obtained in *E. coli* DH5α, the plasmid was transformed to the methyltransferase-deficient *E. coli* strain EC135 to improve transformation efficiency in *L. plantarum* WCFS1 [78].

### Transformation of plasmids to *L. plantarum* WCFS1

Transformation of plasmids into *L. plantarum* WCFS1 was performed as described previously [79]. Briefly, to make electrocompetent cells for transformation, 1 mL of an overnight culture grown in MRS broth at 37 °C without shaking was used to inoculate 25 mL of fresh MRS supplemented with 2.5% glycine and was grown at 37 °C without shaking in 50 mL falcon tube until OD_600_ reached 0.6–0.8. Then, cells were washed twice with 5 mL ice-cold MgCl_2_ (10 mM) and twice more with 5 mL ice-cold SacGly solution. Cells were resuspended in 500 μL ice cold SacGly and aliquoted at 60 μL to be used immediately. Plasmid DNA (1 mg suspended in water) and 60 μL of electrocompetent cells were added to a pre-cooled 1-mm electroporation cuvette and transformed with the following conditions: 1.8 kV, 200 U resistance, and 25 mF capacitance. Following electroporation, cells were recovered in MRS broth for 3 hours at 37 °C and then plated on MRS agar containing appropriate antibiotics for 2-3 days. Chloramphenicol and erythromycin concentrations were both 10 mg/mL in MRS liquid and solid medium.

### Genome editing

To delete the *mprF* gene, electrocompetent *L. plantarum* WCFS1 cells were transformed with pAA032 (recombineering template). Transformants were plated on MRS agar plates containing chloramphenicol. *L. plantarum* WCFS1 harboring the pAA032 were made electrocompetent again and transformed with pAA009 (containing Cas9 and the genome-targeting sgRNA). Transformants were plated on MRS agar containing erythromycin and chloramphenicol for the selection of pAA009 and pAA032. Surviving colonies were screened for the desired genomic deletion using colony PCR with primers oAA036-037, and the PCR products were subjected to gel electrophoresis, PCR clean-up (Macherey-Nagel Cat. No. 740609.250S), and Sanger sequencing (Microsynth GmbH) with primers oAA047-130, which attach to the genome of *L. plantarum* WCFS1 outside of the homology arms to validate the deletion of *mprF* (Supplementary Figure 2). Both plasmids were cured from the mutant *L. plantarum* WCFS1 Δ*mprF* strain by performing a cycle of culturing it in non-selective MRS liquid medium and plating on non-selective MRS solid medium. Then, after each round of non-selective growth, cultures were plated on MRS agar supplemented with either chloramphenicol or erythromycin. This cycle was repeated until the mutant strain was sensitive to both antibiotics.

### Quantification of pathogen load

Flies were infected with bacteria at the indicated OD as described above and allowed to recover. At the indicated time points post-infection, flies were anesthetized using CO2 and surface sterilized by washing them in 70% ethanol. Flies were homogenized using a Precellys TM bead beater at 6,500 rpm for 30 s in LB broth, with 200 μL for pools of 5 flies. These homogenates were serially diluted and plated on LB agar. Bacterial plates were incubated overnight, and colony-forming units (CFUs) were counted manually.

Female flies were used to perform CFU record and gene expression assays. Survival tests were always performed in male flies.

### Generation of germ-free flies

Embryos laid by females over a 12 hours period on grape juice plates were rinsed in 1x PBS and transferred to 1.5 mL Eppendorf tube. All subsequent steps were performed in a sterile hood. After embryos sedimented to the bottom of the tube, PBS was removed and 3% sodium hypochlorite solution was added. After 10 min, the bleach was discarded, and dechorionated embryos were rinsed three times in sterile PBS followed by one wash with 70% ethanol. Embryos were transferred by pipette to tubes with antibiotics-supplemented food and kept at 25 °C. Emerged germ-free adult flies were used for subsequent experiments.

### Estimating *L. plantarum* load in *Drosophila* after infection, priming, and genetic immune activation

We performed colonization and priming following previously published protocol (Arias-Rojas 2023). Briefly, germ-free flies were (a) mono-colonized with either *L. plantarum* wild-type or Δ*mprF* for 48 hours. After the colonization, infection with *Ecc15* was perform and *L. plantarum* load was estimated by plating fly homogenates on MRS agar plates. (b) Germ-free flies were primed with *Ecc15* for 3 hours and mono-colonized with either *L. plantarum* wild-type or Δ*mprF.* CFUs were recorded in by plating fly homogenates on MRS plates. (c) Germ-free flies with overactivated Imd pathway and their controls were monocolonized with *L. plantarum* wild-type or Δ*mprF.* and kept at 29 °C till the CFUs recording. Flies were flipped into conventional vials 48 hours post-treatment. *L. plantarum* was used at OD_600_=50 and *Ecc15* at OD_600_=200. Bacteria were mixed 1:1 with 5% sucrose. 2.5% sucrose was use as control treatment in the infection or priming. For all the mixtures, 150 µL were placed onto paper filter disks covering the fly food surface. Once the solution was absorbed, flies were flipped to these vials for colonization or infection.

### RNA extraction and RT-qPCR

In systemic infection 5 flies per sample were collected at indicated time points. For colonization or oral infection 20 guts were collected at indicated time points. Total RNA was isolated using TRIzol reagent according with the manufactures protocol. After quantification in NanoDrop ND-1000 spectrophotometer, 500 ng of total RNA were used to perform cDNA synthesis, using PrimeScript RT (TAKARA) and random hexamers. qPCR was performed on LightCycler 480 (Roche) in 384-well plates using SYBR Select Master Mix from Applied Biosystems. Expression values were normalized to RP49. Primer sequences are listed in the Supplementary Table S4.

### Lipid extraction

Bacterial cultures were growth from OD_600_=0.1 till they reached OD_600_=5. Cultures were pelleted in 2 mL Eppendorf tubes for 5 min at max speed at room temperature. Pellets were resuspended in 1 mL of 0.9% NaCl. The cells were pelleted again by centrifugation for 5 min at max speed. 5 µL of internal standard was added to each pellet followed by the addition of 120 µL of H_2_O, 150 µL of chloroform and 300 µL of MeOH. The mixture was incubated 10 min on cold shaker at 4 °C. 150 µL of chloroform and 150 µL 0.85% KCl in H_2_O were added after the incubation. The biomass was separated by centrifugation for 5 min at max speed. The lower phase was harvest using a glass inlet. The isolated phase was dried under a nitrogen stream and stored at –20°C.

The relative quantification and lipid annotation were performed by using HRES-LC-MS/MS. The chromatographic separation was performed using an Acquity Premier CSH C18 column (2.1 × 100 mm, 1.7 μm particle size, Water) with a constant flow rate of 0.3 mL/min with mobile phase A being 10 mM ammonium formate in 6:4 acetonitrile:water and phase B being 9:1 isopropanol:acetonitrile (Honeywell, Morristown, New Jersey, USA) at 40 °C. The injection volume was 5 µL. The mobile phase profile consisted of the following steps and linear gradients: 0 – 5 min constant at 5% B; 5 – 20 min from 5 to 98% B; 20 – 27 min constant at 98% B; 27 – 27.1 min from 98 to 5% B; 27.1 – 30 min constant at 5% B. For the measurement, a Thermo Fischer Scientific ID-X Orbitrap mass spectrometer was used. Ionization was performed using a high-temperature electrospray ion source at a static spray voltage of 3,500 V (positive) and a static spray voltage of 2,800 V (negative), sheath gas at 50 (Arb), auxiliary gas at 10 (Arb), and ion transfer tube and vaporizer at 325 °C and 300 °C, respectively.

Data-dependent MS^2^ measurements were conducted by applying an orbitrap mass resolution of 120,000 using quadrupole isolation in a mass range of 200-2,000 and combining it with a high energy collision dissociation (HCD). HCD was performed on the ten most abundant ions per scan with a relative collision energy of 25%. Fragments were detected using the orbitrap mass analyzer at a predefined mass resolution of 15,000. Dynamic exclusion with an exclusion duration of 5 seconds after 1 scan with a mass tolerance of 10 ppm was used to increase coverage.

Due to database limitations in annotating the Lysyl-PG lipids, we used a lipid standard (18:1 Lysyl-PG, Avanti) to identify the fragmentation pattern and possible unique fragments. We found two individual fragments that are lipid class-specific, and with the help of these two fragments, we identified all the Lysyl-PG species present in our measurement. The two fragments are 301.1159 [M+H]^+^ for the positive mode and 145.0982 [M-H]^-^ for the negative mode. Compound Discoverer v3.3.2.31 (Thermo Fisher Scientific) was used to annotate the Lysyl-PG lipids in the sample. We added the two unique fragments in the “Compound Classes” library and applied a workflow seeking for MS/MS spectra in which the two fragments are present. Skyline v22.2.0.255 (MacCoss Lab, University of Washington) was used to get the relative abundance of the different annotated Lysyl-PG species, and the normalized area was extracted and used for further analysis and plotting. The data were normalized by the total ion current defined by Skyline software.

### Antibiotic Inhibition Assay (AIA)

Overnight bacterial cultures were adjusted to OD_600_=0.1, and 50 µL of culture were pipetted into flat bottom 96 well plates prefilled with ranging dilutions of either Polymyxin B (Fischer Scientific), Gentamicin (Sigma), Defensin (Alfa Aesar) or Cecropin B (Sigma). After 6h incubation, bacterial growth was recorded to determine antibiotic inhibition values. Reads were performed in Infinite 200 Pro plate reader (Tecan).

To determine the kinetics of bacterial growth in presence of antimicrobial *in vitro* (Fig. S3), overnight cultures were adjusted to OD_600_=0.05 and grew 20 hours in 96 well plates in a plate reader at 37 °C in MRS medium supplemented with tested antibiotics. Bacterial growth was estimated by measuring OD_600_ in Infinite 200 Pro plate reader (Tecan).

### Cytochrome C and 5-FAM-LC-LL37 binding

Bacterial cultures were grown to OD_600_=0.6, washed once with 1x PBS and resuspended in Buffer A (1 M KH_2_PO_4_, pH 7.0, BSA 0.01%) and Cytochrome C (Sigma) solution (0.5 mg/mL), the cells were incubated 15 min at RT. Supernatants were obtained by centrifugation and measured in 96 well plates at 440 nm. Overnight cultures were adjusted to OD_600_ 0.1 in PBS 1x, and 5-FAM-LC-LL37 (Eurogentec) solution (14 µM) was added to each sample and incubated 1 hour at 37 °C, 590 rpm. Supernatants were obtained by centrifugation and transferred to 96 well plates to measure the Fluorescence (absorbance 494 nm and emission 521 nm). Reads were performed in Infinite 200 Pro plate reader (Tecan).

### Scanning electron microscopy

Bacterial cells were fixed with 2.5% glutaraldehyde and 20 µL drops of bacterial suspension were spotted onto polylysine-coated round glass coverslips place into the cavities of a 24-well cell culture plate. After 1 h of incubation in a moist chamber, PBS was added to each well, and the samples were fixed 2.5% glutaraldehyde for 30 min. Sample were washed and post-fixed using repeated incubations with 1% osmium tetroxide and 1% tannic acid, dehydrated with a graded ethanol series, critical point dried and coated with 3 nm platinum/carbon. Specimens were analysed in a Leo 1550 field emission scanning electron microscope using the in-lens detector at 20 kV. For quantification, images were recorded at a magnification of 2000x and analysed with the Volocity 6.5.1 software package.

### Peptidoglycan release assay and peptidoglycan isolation

Overnight cultures were set to OD_600_=0.1 and grown to OD_600_=2 stationary in MRS media. Lysozyme (10 mg/mL) was added during the OD_600_=0.5. Bacterial cultures were centrifuged and supernatants were heated in the thermoblock at 95 °C for 20 min. 69 nL of supernatants were injected into the thorax of flies. Peptidoglycan isolation was performed as described in (Arias-Rojas et al., 2023). 9.2 nL of isolated purified peptidoglycan was injected into the thorax of the flies. For peptidoglycan feeding, 150 µL of 15 mg/mL of isolated peptidoglycan solution in LAL water (Invivogen) was mixed 1:1 with 5% sucrose and fed to flies from filter discs to test the expression of *Dpt* in the gut upon PNG ingestion. Batches of 20 females flies (10 days old), per sample were use. Either after injection or ingestion flies were kept at 29 °C for 4 hours. Drummond scientific Nanoject II was use to inject flies (Drummond, Broomall, PA).

### Generation of complemented *L. plantatum* Δ*mprF* mutant

For complementation, we used Gram-positive/Gram-negative *shuttle vector* pSIP409 [80] offering inducible expression in *Lactobacilli. mprF* gene was PCR amplified with proof-reading Phusion polymerase (Thermo Fisher Scientific) using genomic DNA of *L. plantarum* WCFS1 as template and mprF NcoI F/ mprF XhoI R primers containing restriction digest sites. PCR product and pSIP409 plasmid were digested with NcoI and XhoI enzymes (NEB), gel purified with *Monarch* DNA *Gel Extraction Kit* (NEB), and ligated with T4 DNA ligase (Thermo Fisher Scientific). Ligation products were transformed into chemically competent TOP10 *E. coli* and positive transformant were selected on LB agar with 150 µg/mL erythromycin. After sequence verification, the obtained plasmid pSIP409-Lp-mprF was electroporated into the *L. plantarum* Δ*mprF* mutant as described above to generate complemented mutant strain *L. plantatum* Δ*mprF pSIP409-Lp-mprF*. The complemented strain was grown overnight stationary in MRS at 37 °C. Next day cultures were diluted to OD_600_=0.1 and grown to OD_600_=0.3. At this OD, MprF expression was induced using 12.5 ng/mL of the peptide pheromone IP-673. Uninduced culture was used as a control. Cultures were grown for another 3 hours, harvested by centrifugation (3,600 rpm for 15 min) and finally adjusted to OD_600_=0.5 before being fed to female flies.

### Generation of *E. coli* expressing *L. plantatum* MprF

For expression in *E. coli*, we cloned the *L. plantarum mprF* gene into pBAD18 expression vector using restriction digest with BamHI/SalI and ligation. The obtained plasmid pBAD18-LpMprF was sequence verified and transformed into *E. coli* TOP10 generating *E. coli-pBAD18-LpMprF* strain. Considering that pBAD18 is an arabinose-inducible vector, *E. coli-pBAD18-LpMprF* grown in LB without arabinose was used as a control, while the same strain raised in the presence of 0.2% of arabinose was studied as an MprF producer.

### LTA analysis by western blot

#### Crude extract preparation

Bacteria were grown stationary in 50 mL MRS at 37 °C overnight. After homogenizing, 20 mL were harvested and pelleted at 3,200 x *g* for 10 min. The pellets were then washed in 200 µL of 50 mM citric buffer (pH 4.7) and the optical density of both solutions was normalized to the same OD_600_. After centrifugation, the pellets were resuspended in 600 µL of solution B (equal volume of citric buffer and 2x LDS Buffer), heated at 90 °C for 30 min, and then cooled on ice for 3 min. The samples were incubated with 100 U of benzonase for 30 min at 37 °C and then centrifuged at 4 °C and 3,000 x *g* for 20 min. After centrifugation, the supernatants were heated at 70 °C for 10 min.

### Western Blot

12 µL of supernatants were separated on a Bolt 4-12% Bis-Tris Plus Gel (Invitrogen) for 25 min at 200 V. After migration, the samples were transferred onto a membrane using an iBlot 2 NC Mini Stack device (Invitrogen) and the following protocol: 1 min at 20 V, 4 min at 23 V, and 2 min at 25 V. The membrane was then blocked on a shaking board at room temperature for one hour using 5% non-fat milk in PBS-T (PBS 1X + 0.1% TWEEN 20). After washing in PBS-T (3 x 5 min), the membrane was incubated in primary antibody solution (Gram-positive LTA monoclonal antibody (MA1-7402, Thermo Fischer Scientific) diluted 1:1000) at 4 °C overnight on a rolling board. After washing in PBS-T (3 x 5 min), the membrane was incubated in secondary antibody (anti-mouse HRP-linked secondary antibody (P0447, Agilent/Dako) diluted 1:10,000) at room temperature for two hours. The membrane was washed in PBS-T (3 x 5 min) one last time before by chemiluminescent detection and imaging. Western blot analysis of purified LTA was performed in the same way, with the exception that defined amounts of purified LTA were separated on a gel.

### LTA purification, de-O-acylation, and analysis by NMR and Tris-tricine-PAGE

LTA isolation and purification were performed as described elsewhere [81]. For de-O-acylation, LTA preparations were dissolved 5 µg/µL in 1 M hydrazine (N_2_H_4_) in THF (Sigma Aldrich, 433632) and 20 µL Millipore-water were added for better solubility. After mixing, samples were incubated for 1 hour at 37 °C under stirring. The reaction was quenched by careful adding of ice-cold acetone (same volume as the N_2_H_4_/THF-solution) and subsequently dried under a stream of nitrogen. The latter step was repeated twice. For desalting, the de-O-acylated LTA was dialyzed against water (MWCO: 500–1000 Da) including three water exchanges and one overnight dialysis.

NMR spectroscopic measurements were performed in D_2_O (purchased from Deutero GmbH (Kastellaun, Germany)) at 300 K on a Bruker Avance^III^ 700 MHz (equipped with an inverse 5 mm quadruple-resonance Z-grad cryoprobe). Acetone was used as an external standard for calibration of ^1^H (δ_H_ = 2.225) and ^13^C (δ_C_ = 30.89) NMR spectra [82] and 85% of phosphoric acid was used as an external standard for calibration of ^31^P NMR spectra (δ_P_ = 0.00). All data were acquired and processed by using Bruker TOPSPIN V 3.0 or higher. ^1^H NMR assignments were confirmed by 2D ^1^H,^1^H-COSY and total correlation spectroscopy (TOCSY) experiments. ^13^C NMR assignments were indicated by 2D ^1^H,^13^C-HSQC, based on the ^1^H NMR assignments. Interresidue connectivity and further evidence for ^13^C assignment were obtained from 2D ^1^H,^13^C-heteronuclear multiple bond correlation and ^1^H,^13^C-HSQC-TOCSY. Connectivity of phosphate groups were assigned by 2D ^1^H,^31^P-HMQC and ^1^H,^31^P-HMQC-TOCSY.

De-O-acylated LTA were subjected to native Tris-tricine-PAGE analysis essentially following a published protocol [63]. First, aliquots of the de-O-acylated LTA were dissolved in Millipore-water in a concentration of 5 µg/µL. Portions of appr. 80 µL were then applied to benzonase and subsequent proteinase K digestion. For this, the respective portion was mixed with an equal volume of a mixture of Millipore-water/100 mM Tris-HCl (pH 8.0)/20 mM MgCl_2_/benzonase (25 U/µL) 0.8/1.0/0.5/0.2 (v/v/v/v). The 25 U/µL benzonase solution was freshly prepared by mixing the commercial 250 U/µL benzonase solution (1.01695.0001, Merck) with 100 mM Tris-HCl (pH 8.0)/20 mM MgCl_2_/Millipore-water in a 1:2:1:6 (v/v/v/v) ratio. After an incubation for 2 hours at 37 °C, a proteinase K solution (20 mg/mL; AM2548, Ambion) was added in a volume equivalent to 1/32 of this mixture and the resulting mixture further incubated for 2 hours at 50 °C. The final solutions of such enzymatic digests have an LTA concentration of 2.42 mg/mL. The treated samples were stored at –20 °C until they were applied to Tris-tricine PAGE. 15 µg material in 25 µL solution [7.4 µL enzymatic digest, 15.1 µL Millipore-water, 7.5 µL 4x loading dye (according to [63])] were loaded on the PAGE. Electrophoresis was performed at 14 mA (gel dimension: 16 cm x 14 cm x 0.75 mm) and 4 °C for 877 min in a Hoefer SE600 Gel Electrophoresis Unit (Hoefer Inc., Holliston, MA, USA). Subsequent sequential alcian blue [63] and silver staining [83] were performed as described, respectively.

### Statistical analysis

Statistical test was conducted using R version 4.3.2. Survival analysis was carried with the Kaplan–Meier method, and the Log Rank test, using R package survminer. Statistical parameters and tests are shown in the figure legends. The interquartile range from the first to third quartiles, with whiskers representing the tenth and ninetieth percentiles are shown in the dot plots. Pairwise comparisons were executed and plotted collectively. Data visualization was performed with the R packages ggplot2, dplyr, reshape2, and tidyverse.

## Availability of data and materials

All data generated or analysed during this study are included in this published article and its supplementary information files.

## Acknowledgements

We are grateful to Andreas Peschel (Eberhard Karl University of Tübingen) for kindly providing the *S. aureus* Δ*mprF* mutant. We thank Pascal Hols (Université catholique de Louvain) for sharing the pSIP409 plasmid and Simone Thomsen, Madleen Reddig and Heiko Käßner (all RC Borstel) for excellent technical assistance.

## Funding

This work was supported by the Max Planck Society. Funding from the Deutsche Forschungsgemeinschaft grant is acknowledged by I.I. (IA 81/2-1) and N.G. (GI 979/1-2) and the European Research Council Consolidator Grant is acknowledged by C.L.B. (865973). Open Access funding was enabled and organized by Projekt DEAL. The funders had no role in study design, data collection, and interpretation, or the decision to submit the work for publication.

## Contributions

II and AAR conceived and designed the study. AAR, AA, GA, BA, US, EF, DF, VB, NP, NG conducted the experiments. AAR, AA, GA, NG, CLB, NP, II wrote the manuscript. AAR, AA, GA, VB, NP, NG analysed and interpreted the data. All authors reviewed and approved the final manuscript.

## Ethics declarations

### Ethics approval and consent to participate

Not applicable

### Consent for publication

Not applicable

### Competing interests

The authors declare no competing interests.

## Notes

### Competing Interest Statement

The authors have declared no competing interest.

