## Supplementary information for "MprF-mediated immune evasion is necessary for *Lactiplantibacillus plantarum* resilience in *Drosophila* gut during inflammation"

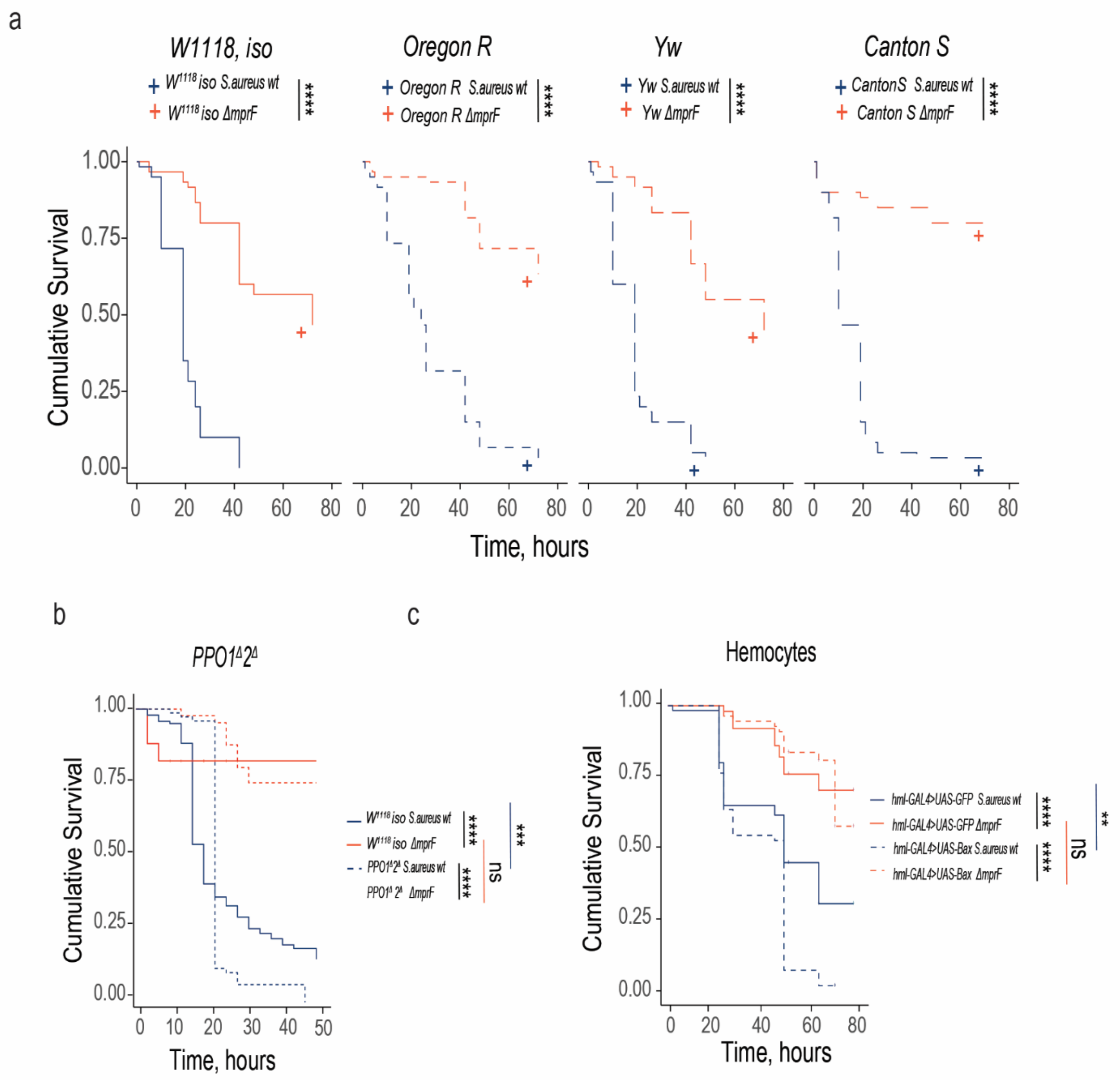


**Figure S1. Melanization and hemocytes are not involved in the control of *Staphylococcus aureus* Δ*mprF* mutant.** (**a**) Survival rates of *Drosophila* wild-type strains infected with wild-type *S. aureus* or *S. aureus* Δ*mprF* mutant. (**b, c**) Survival rates of melanisation-deficient mutant (*PPO1*$^{\Delta}$*2*$^{\Delta}$) (**b**) and hemocytes-depleted flies (**c**) infected with wild-type *S. aureus* or *S. aureus* Δ*mprF* mutant. Each survival graph shows cumulative results of three independent experiments.

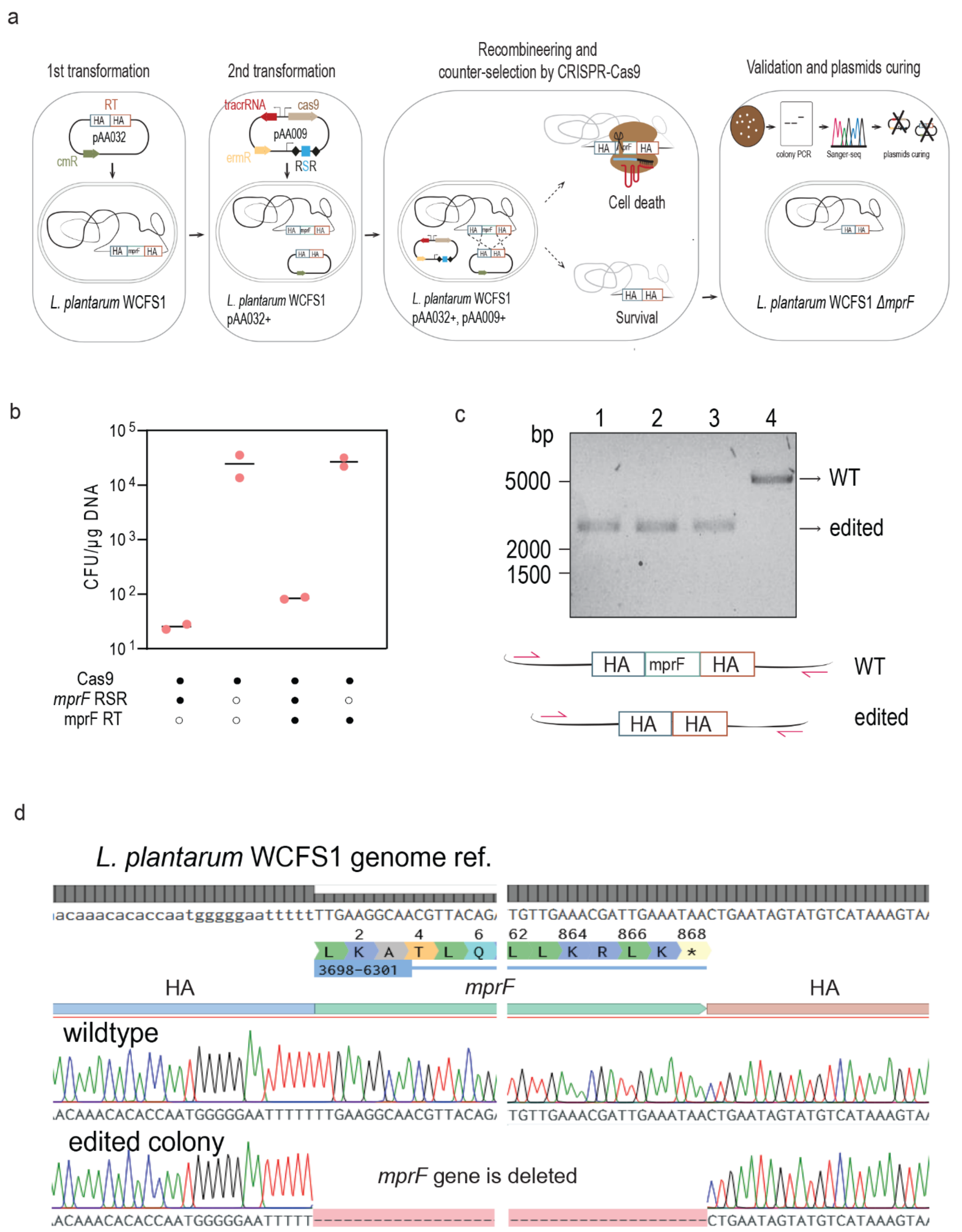


**Figure S2. Generation of *L. plantarum* Δ*mprF* mutant.** (**a**) Scheme illustrating CRISPR-Cas9 genome editing approach to knock out *mprF* in *L. plantarum* WCFS1. First, a recombineering template (RT) plasmid pAA032 containing approximately 250-bp homology arms (HA9) flanking *mprF* gene was transformed into *L. plantarum*. Then, the shuttle vector pAA009 encoding SpyCas9, its tracrRNA, and single-spacer CRISPR array targeting a site within *mprF* was transformed into the *L. plantarum* containing recombineering template plasmid to counter-select the unedited cells. Surviving colonies were screened by colony PCR (cPCR) that amplifies the genome of *L. plantarum*, but not the plasmid with the recombineering template. (**b**) Killing activity of the guide RNA containing *mprF* targeting spacer. The observed 1,000-fold reduction in colony forming units (CFU) in the presence of targeting repeat-spacer-repeat (RSR) indicated targeting and cleaving activities on the genome containing *mprF* gene (**c-d**) Validation of *mprF* deletion via colony PCR (**c**) and Sanger sequencing (**d**). cPCR was performed using primers that bind to the genome but not the plasmid. PCR products of ~2,400 bp indicated clean deletion of the ~2,600-bp *mprF* gene. PCR products were then subjected to Sanger sequencing to confirm that *mprF* gene was successfully deleted from the start codon to the stop codon. After confirmation, plasmids were cured to generate the final strain.

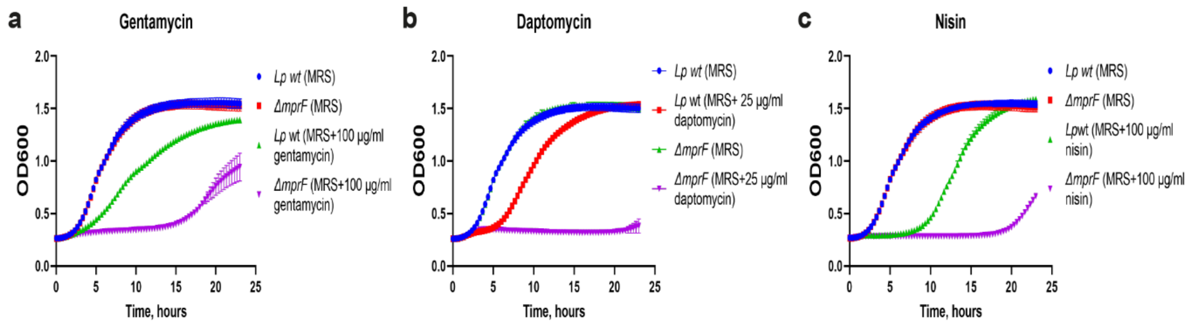


**Figure S3. Susceptibility to antibiotics. (a-c)** Growth of *L. plantarum* wild-type and *L. plantarum* Δ*mprF* mutant in MRS media (control) and MRS media supplemented with 100 µg/ml gentamicin (**a**), 25 µg/ml daptomycin (**b**), and 100 µg/ml nisin (**c**). Data show mean and SD of 3 independent experiments.

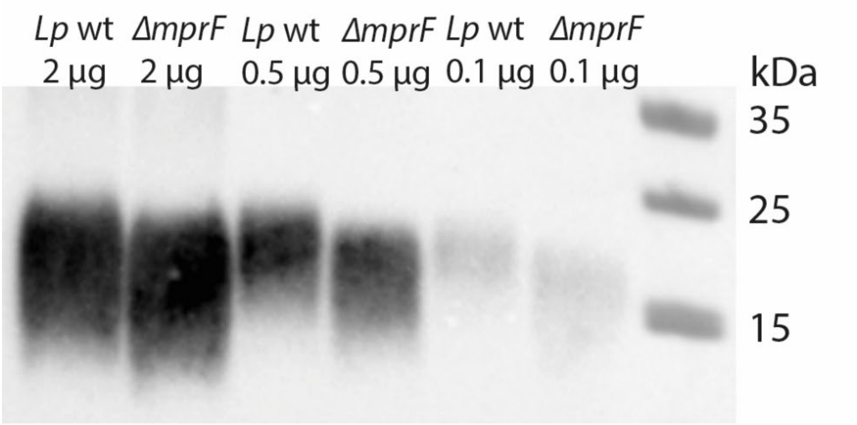


**Figure S4. Differences in the profile of purified LTA**. LTA profile of *L. plantarum* wild-type and Δ*mprF* mutant detected with Western blot and anti-LTA MAb at a 1:1000 dilution. LTA was purified from *L. plantarum* wild-type and Δ*mprF* mutant. Indicated amounts of purified LTA were mixed with 2X LDS Buffer and resolved on a Bolt 4-12% Bis-Tris Plus Gel. Subsequent Western blot was performed and showed reduced size of LTA from Δ*mprF* mutant.

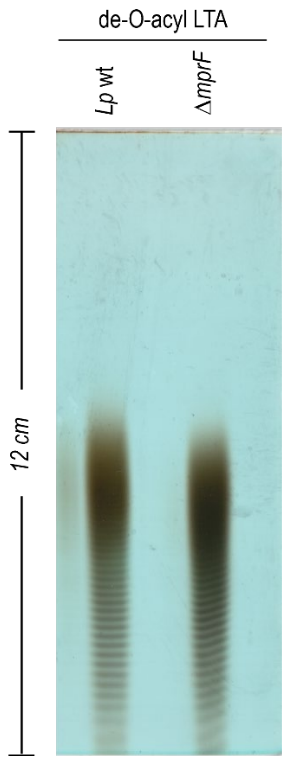


**Figure S5. Full length separation gel of the Tris-tricine-PAGE with combined alcian blue and silver staining of de-O-acylated LTA.** Truncated version of this gel is depicted in Fig. 6b.

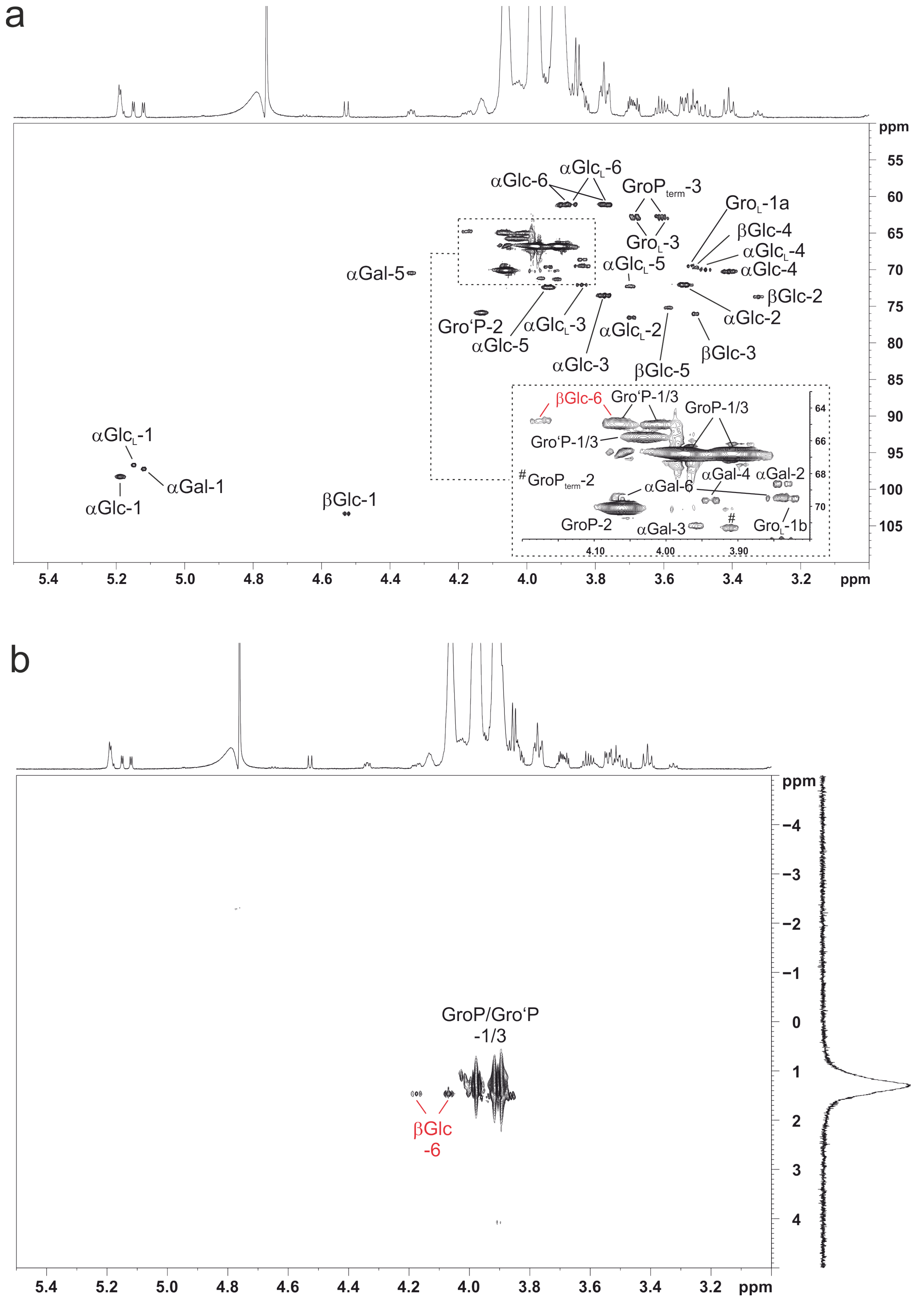


**Figure S6. NMR analysis of de-O-acylated LTA of *L. plantarum* WCFS1 reveals that the poly-glycerolphosphate chain is attached to the O-6 position of the βGlc$_p$ residue. (a)**

Shown is a section ($\delta_H$ 5.5-3.0 ppm; $\delta_C$ 110-50 ppm) of the $^1H,^{13}C$-HSQC NMR spectrum (recorded in $D_2O$ at 300 K as dept-version) including signal assignment. (**b**) Shown is a section ($\delta_H$ 5.5-3.0 ppm; $\delta_P$ 5-(–5) ppm) of the $^1H,^{31}P$-HMQC NMR spectrum (recorded in $D_2O$ at 300 K) including signal assignment. In both panels, the cross-correlations for the O-6 position of βGlc*p* are highlighted in red. The corresponding NMR chemical shift data are listed in Table S1.

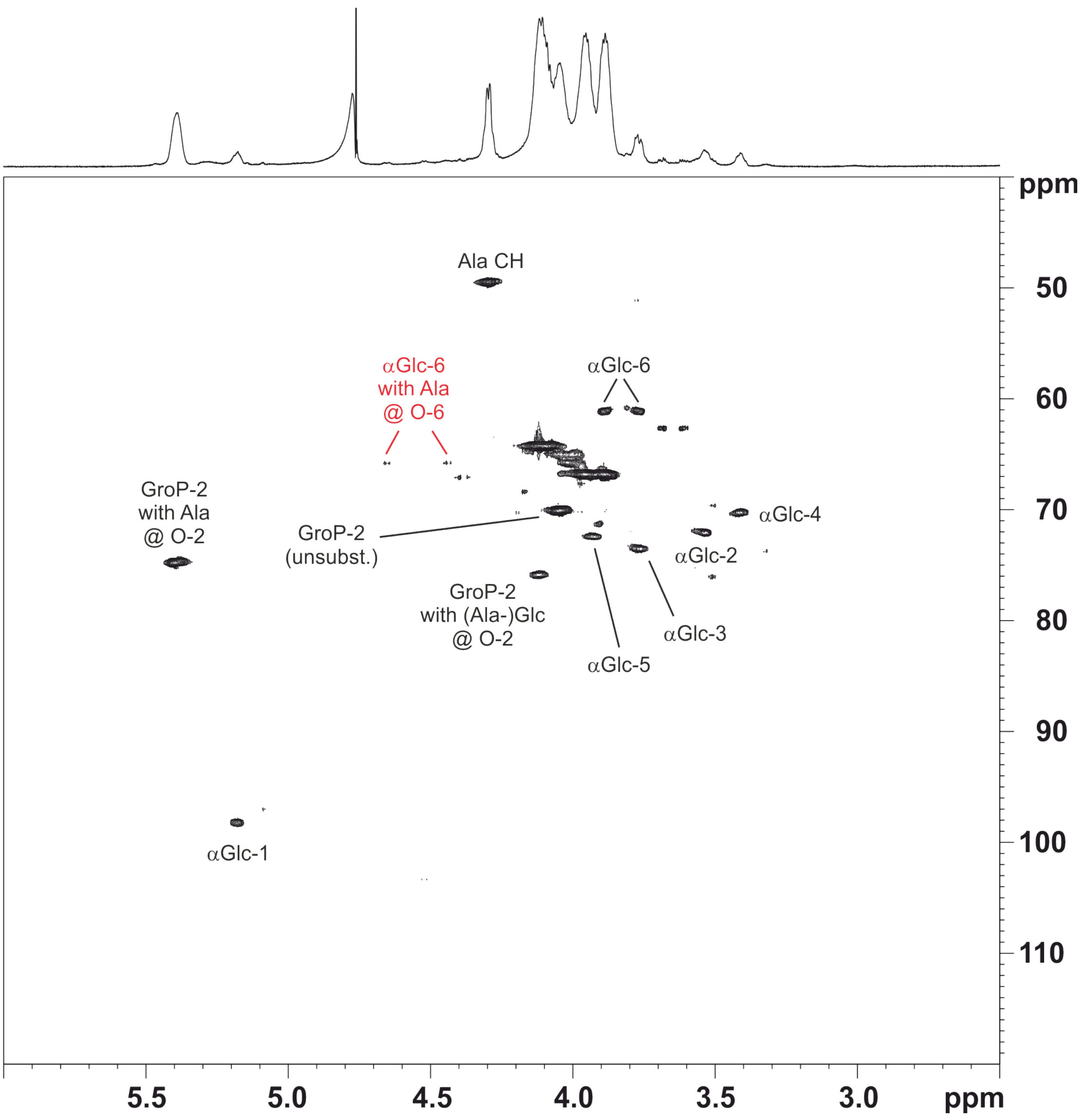


**Figure S7**. ***L. plantarum* WCFS1 LTA contains alanine (Ala) and αGlc*p* residues as major substituents at the O-2 position of the glycerolphosphate (GroP) moieties. In addition, there is evidence for a small proportion of 6-Ala-αGlc*p*** (as described for *Lp* strain NC8 [1]. Shown is a section ($\delta_H$ 6.0-2.5 ppm; $\delta_C$ 120-40 ppm) of the $^1H,^{13}C$-HSQC NMR spectrum (recorded in $D_2O$ at 300 K as dept-version) obtained from native LTA of the wild-type strain.

**Table S1. $^{1}$H (700.4 MHz), $^{13}$C NMR (176.1 MHz), and $^{31}$P NMR (283.5 MHz) chemical shift data (δ, ppm) [*J*, Hz] for *L. plantarum* strain WCFS1 LTA after hydrazine treatment (de-O-acyl LTA) recorded in $D_2O$ at 300 K.**

| **Residue**<br>**[assignment]** | **H-1**<br>***C-1*** | **H-2**<br>***C-2*** | **H-3**<br>***C-3*** | **H-4**<br>***C-4*** | **H-5**<br>***C-5*** | **H-6**<br>***C-6*** |
|---|---|---|---|---|---|---|
| Gro-(1→ [**Gro$_{L}$**] | 3.84-3.81*<br>3.53.3.50*<br>*69.6* | *n.d.*<br><br>*n.d.* | 3.70-3.67*<br>3.63-3.59*<br>*62.8* | | | |
| →1)-α-Glc*p*-(2→ [α**Glc$_{L}$**] | 5.15 [3.5]<br><br>*96.8* | 3.71-3.68*<br><br>*76.6* | 3.86-3.82*<br><br>*72.2* | 3.50-3.46*<br><br>*70.2* | 3.72-3.68*<br><br>*72.4* | 3.89-3.85*<br>3.79-3.75*<br>*61.2* |
| →1)-α-Gal*p*-(6→ [α**Gal**] | 5.12 [4.1]<br><br>*97.3* | 3.86-3.82*<br><br>*68.8* | 3.97-3.94*<br><br>*71.3* | 3.95-3.93*<br><br>*69.7* | 4.36-4.32*<br><br>*70.6* | 4.08-4.04*<br>3.87-3.82*<br>*69.6* |
| →1)-β-Glc*p*-(6→*P* [βG**lc**] | 4.53 [8.0]<br><br>*103.4* | 3.34-3.30*<br><br>*73.8* | 3.52-3.49*<br><br>*76.1* | 3.52-3.49*<br><br>*69.8* | 3.60-3.56*<br><br>*75.3* | 4.20-4.16*<br>4.09-4.05*<br>*65.0* |
| *P*→1)-Gro-(3→ [**GroP**] | 4.01-3.95*<br>3.94-3.88*<br>*66.9 [5.2]* | 4.09-4.03*<br><br>*70.2* | 4.01-3.95*<br>3.94-3.88*<br>*66.9 [5.2]* | | | |
| *P*→1, αGlc→2)-Gro-(3→ [**Gro'P**][$] | 4.06-4.00*<br>*65.9* | 4.16-4.11*<br>*76.0* | 4.06-4.00*<br>*65.9* | | | |
| | | and | | | | |
| | 4.09-4.04*<br>4.04-3.99*<br>*65.1* | 4.16-4.11*<br><br>*76.0* | 4.09-4.04*<br>4.04-3.99*<br>*65.1* | | | |
| *P*→1)-Gro [**Gro$^{term}$**] | 3.96-3.92*<br>3.89-3.85*<br>*67.0* | 3.93-3.90*<br><br>*71.4* | 3.70-3.67*<br>3.63-3.59*<br>*62.7* | | | |
| α-Glc*p*-(1→ [α**Glc**] major | 5.19 [3.6]<br><br>*98.3* | 3.54<br>[10.3, 3.6]<br>*72.2* | 3.80-3.75*<br><br>*73.6* | 3.41 [9.6,<br>9.5]<br>*70.3* | 3.96-3.92*<br><br>*72.5* | 3.92-3.87*<br>3.79-3.75*<br>*61.2* |
| minor | 5.18 [3.6]<br>*98.4* | 3.54-3.51*<br>*72.2* | | | | |
| **$^{31}$P** | Gro-*P*-Gro$_{term}$ 1.52[#]; βG**lc**-6-*P*-Gro 1.48[#]; Gro-*P*-Gro / Gro-*P*-Gro‘ / Gro‘-*P*-Gro‘ 1.70-0.95 | | | | | |

*non-resolved multiplet; [#]values determined using $^{1}$H,$^{31}$P-HMQC-TOCSY; [$]two different signals for Gro'P-1/3 are present due to varying nature of the neighboring repeating units (possible are the combinations 2 x GroP, 1 x GroP + 1 x Gro'P or 2 x Gro'P); *n.d.* = not detected.

**Table S2. Bacterial strains used in this study.**

| Strain | Growth conditions | Relevant characteristics | Reference or sourse |
|---|---|---|---|
| *Pectobacterium carotovorum carotovorum (Ecc15)* | 29°C Shaking incubator, in LB media | Natural *Drosophila* pathogen | Basset et al., 2000 [2] |
| *Lactiplantibacillus plantarum NCIMB 8826 (WCFS1)* | 37°C Stationary Incubator, in MRS media | Strain with high transformation efficiency | Kleerebeyem et al 2003 [3] |
| *Lactiplantibacillus plantarum ∆mprF* | 37°C Stationary Incubator, in MRS media | NCIMB strain deleted for mprF gene | This study |
| *L. plantatum ∆mprF pSIP409-Lp-mprF* | 37°C Stationary Incubator, in MRS media supplemented with IP-673 peptide | NCIMB strain deleted for mprF gene, containing pSIP409::mprF plasmid for MprF overexpression | This study |
| *Staphylococcus aureus 113* | 37°C Shaking incubator, in TSB media | Pathogen use for Drosophila systemic infection | Peschel et al., 2001 [4] |
| *Staphylococcus aureus ∆mprF* | 37°C Shaking incubator, in TSB media | S.aureus 113 strain deleted for MPRF genes | Peschel et al., 2001 [4] |
| *E. coli TOP10* | 37°C Shaking incubator, in LB media | F–mcrA Δ(mrr-hsdRMS-mcrBC) φ80lacZΔM15 ΔlacX74 recA1 araD139 Δ(ara-leu)7697 galU galK λ–rpsL(StrR) endA1 nupG | Thermo Fisher Scientific |
| *E. coli-pBAD18-LpMprF* | 37°C Shaking incubator, in LB media suplemented with Ampicillin and L-arabinose 0.2% | E. coli TOP10 strain, containing pBAD18 plasmid for LpMprF overexpression | This study |
| *E. coli DH5a* | 37°C Shaking incubator, in LB media | *F– φ80lacZΔ M15 Δ (lacZYA-argF) U169 recA1 endA1 hsdR17 (rK– mK+) phoA supE44 λ- thi–1 gyrA96 relA1* | Thermo Fisher Scientific |
| *E. coli EC135* | 37°C Shaking incubator, in LB media | *recA1 recA+ mcrA Δ(mrr-hsdRMS-mcrBC) Δdcm::FRT Δdam::FRT* | Zhang et al., 2012 [5] |

### Table S3. Plasmids used in this study.

| CBS ID | Plasmid ID | Description | Plasmid map | Source | Use | Resistance |
|---|---|---|---|---|---|---|
| CBS-4146 | pCB591 | Backbone for RT plasmid | https://benchling.com/s/seq-EZ0vHyRUqOoWOlH3zF4P?m=slm-Il8mCsgv7yZSqDWPZHxF | Leenay et al., 2019 [6] | Cloning | Amp100/Cm10 |
| CBS-4100 | pCB578 | Cas9 + tracrRNA + *ackA* RSR backbone for targeting plasmid | https://benchling.com/s/seq-CE1TOCpvMS7aVMML8oUC?m=slm-e6Ha9sWlj5BClVFEERtN | Leenay et al., 2019 [6] | Cloning | Erm300/Erm10 |
| CBS-4101 | pCB577 | Cas9 + tracrRNA without RSR | https://benchling.com/s/seq-9KP7wflTR5qRk42gaxba?m=slm-Pz6BgawZrpk50JCTtDZZ | Leenay et al., 2019 [6] | Fig.S2 | Erm300/Erm10 |
| CBS-4148 | pAA032 | Final RT for clean deletion *mprF* in *L.plantarum* WCFS1_250-bp upstream HA + 250-bp downstream HA | https://benchling.com/s/seq-ahIHCDXtqhxqzi6OdnRW?m=slm-hlx6hw9Mzes5ovINbI1W | This study | Fig.S2 | Amp100/Cm10 |
| CBS-3446 | pAA009 | SpyCas9 + tracrRNA + RSR for *mprF* *L.plantarum* WCFS1 | https://benchling.com/s/seq-m6uWIM3tLRhdvmf6GbEg?m=slm-QCQffVNSbwwVnAgIjiSM | This study | Fig.S2 | Erm300/Erm10 |

**Table S4. Primers used in this study.**

| Primer name | Reference or description | Sequence (5‘ – 3‘) |
|---|---|---|
| Defensin F | This study | AGTTCTTCGTTCTCGTGGCT |
| Defensin R | This study | CCACATCGGAAACTGGCTGA |
| Drosomycin F | Dudzic et al 2019 [7] | CGTGAG AACCTTTTCCAATATGAT |
| Drosomycin R | Dudzic et al 2019 [7] | TCCCAGGACCACCAGCAT |
| RP49 F | Iatsenko et al 2018 [8] | GACGCTTCAAGGGACAGTATCTG |
| RP49 R | Iatsenko et al 2018 [8] | AAACGCGGTTCTGCATGAG |
| DptA F | Iatsenko et al 2018 [8] | TGGTGGAGTGGGCTTCAT |
| DptA R | Iatsenko et al 2018 [8] | GCTGCGCAATCGCTTCTA |
| PGRP-LB F | This study | GGCATGATTTACACCGGCAG |
| PGRP-LB R | This study | TCTCCGATCAGCACAATGCC |
| Pirk F | This study | GCTGCAATGGACTGCTCAAG |
| Pirk R | This study | AGAGCTGGGCCTTTTCTTGG |
| oAA027 | Oligo for inserting *mprF* target spacer FWD | CGACGGAAGTCGGCAAAATTGCGAGTTTTAGAGCTATGCTGTTTTGA ATGGTCCCAAAACATGC |
| oAA028 | Oligo for inserting *mprF* target spacer REV | GGCCGCATGTTTTGGGACCATTCAAAACAGCATAGCTCTAAAACTCG CAATTTTGCCGACTTCCGTCGAT |
| oAA097 | Primer for amplifying the genome region of WCFS1 containing *mprF* gene and 250-bp upstream and downstream FWD | ACATAAACGGTAAAGGTTGGTAAAGC |
| oAA098 | Primer for amplifying the genome region of WCFS1 containing *mprF* gene and 250-bp upstream and downstream REV | AGGCTAACCTCGACCTATTC |
| oAA094 | Primer for amplifying the backbone for RT plasmid FWD | GAATAGGTCGAGGTTAGCCTGCTCAAGCTTTCTTTGAACC |
| Pirk R | This study | AGAGCTGGGCCTTTTCTTGG |
| oAA099 | Primer for amplifying the backbone for RT plasmid REV | CCAACCTTTACCGTTTATGTAACTAGTGGTACCTTAGCTGTCAGC |
| oAA033 | Primer to remove *mprF* from recombineering template plasmid FWD | CTGAATAGTATGTCATAAAGTAAG |
| oAA034 | Primer to remove *mprF* from recombineering template plasmid REV | AAAAATTCCCCCATTGGTG |
| oAA038 | Primer on the plasmid backbone for amplifying the RT region for colony PCR FWD | TTTTGCTCACATGTTCTTTC |
| oAA039 | Primer on the plasmid backbone for amplifying the RT region for colony PCR REV | CTGCTTTTTGGCTATCAATC |
| oAA036 | Primer on genome for cPCR of mutant FWD | CGTCAGTTGCTTGTCATTAT |
| oAA037 | Primer on genome for cPCR of mutant REV | AAAGACCGTTCTGAAAAGCA |
| oAA047 | Primer on genome for sequencing FWD | CGAGTTTTTTTGCATGGTGCATAAG |
| oAA130 | Primer on genome for sequencing REV | ACGACGATCTAGTCGCCATG |
| mprF F BamHI | Forward primer for *mprF* cloning into pBAD18 plasmid | CGCGGATCCGATGAAGGCAACGTTACAGAAA |
| mprF R SalI | Reverse primer for *mprF* cloning into pBAD18 plasmid | ACGCGTCGACTTATTTCAATCGTTTCAACAACCAT |
| mprF NcoI F | Forward primer for *mprF* cloning into pSIP409 plasmid | TAAGCACCATGGatgaaggcaacgttacagaa |

| | | |
|---|---|---|
| mprF XhoI R | Reverse primer for *mprF* cloning into pSIP409 plasmid | TAAGCACTCGAGttatttcaatcgtttcaaca |
